# Sex and species associated differences in Complement-mediated immunity in Humans and Rhesus macaques

**DOI:** 10.1101/2023.10.23.563614

**Authors:** Natasha S. Kelkar, Benjamin S. Goldberg, Jérémy Dufloo, Timothée Bruel, Olivier Schwartz, Ann J. Hessell, Margaret E. Ackerman

## Abstract

The complement system can be viewed as a ‘moderator’ of innate immunity, ‘instructor’ of humoral immunity, and ‘regulator’ of adaptive immunity. While sex and aging are known to affect humoral and cellular immune systems, their impact on the complement pathway in humans and rhesus macaques, a commonly used non-human primate model system, have not been well-studied. To address this knowledge gap, we analyzed serum samples from 90 humans and 75 rhesus macaques for the abundance and activity of the complement system components. While sequences of cascade proteins were highly conserved, dramatically different levels were observed between species. Whereas the low levels detected in rhesus samples raised questions about the suitability of the test, differences in levels of complement proteins were observed in male and female humans. Levels of total and antibody-dependent deposition of C1q and C3b on a glycosylated antigen differed between human and rhesus, suggesting differential recognition of glycans. Functional differences in complement-mediated lysis of antibody-sensitized cells were observed in multiple assays and showed that human females frequently exhibited higher lytic activity than human males or rhesus macaques, which typically did not exhibit such sexual dimorphism. Other differences between species and sexes were observed in more narrow contexts—for only certain antibodies, antigens, or assays. Collectively, these results expand our knowledge of sexual dimorphism in the complement system in humans, identifying differences that appear to be absent from rhesus macaques.

## Introduction

Biological factors both influence the composition and development of the immune system and its responses to pathogens. It is believed that sex-based differences in immunity are a consequence of genetic differences attributable to the X chromosome, which encodes immunity genes such as Toll-like receptors, cytokine receptors, genes involved in B cell and T cell activity, transcriptional and regulatory factors(Fish, 2008). Conversely, the Y chromosome, which is exclusively present in males, encodes for genes involved in inflammatory pathways (Flanagan, 2014). Hormones further contribute to the difference between male and female immune responses (Fish, 2008).

Sexual dimorphism in immune responses evolved in diverse species ranging from insects, lizards, birds and mammals (Klein & Flanagan, 2016). For example, many genes that encode for innate immune signaling proteins in *Drosophila melanogaster* are found on the X chromosome and show sex-specific induction in bacterial and fungal infection (Hill-Burns & Clark, 2009, Taylor & Kimbrell, 2007). In humans, sex-based differences have been demonstrated in infectious diseases like COVID-19 (Bienvenu, Noonan et al., 2020, Gadi, Wu et al., 2020, Gersh, O’Keefe et al., 2021, Qi, Ngwa et al., 2021, Zhao, Xu et al., 2021), HIV (Collazos, Asensi et al., 2007), influenza (Wang, Lashua et al., 2022), and mumps (Riggenbach, Haralambieva et al., 2022). Both flow cytometric and single-cell transcriptomics experiments have revealed that females have a lower percentage of natural killer cells in peripheral blood as compared to males (Abdullah, Chai et al., 2012, Huang, Chen et al., 2021). Studies also demonstrate that females have higher phagocytic activity of macrophages and neutrophils (Spitzer, 1999), and higher CD4/CD8 ratios as compared to age-matched males (Abdullah et al., 2012, Amadori, Zamarchi et al., 1995, Lee, Yap et al., 1996, Lisse, Aaby et al., 1997, Uppal, Verma et al., 2003, Wikby, Mansson et al., 2008) and more efficient antigen presentation than in males (Weinstein, Ran et al., 1984). Sex-based differences in vaccine-induced humoral immunity have been seen in children and adults. Adult females are generally known to develop higher antibody titers due to enhanced immune activation than their male counterparts (Fischinger, Boudreau et al., 2019, Klein, Jedlicka et al., 2010), a difference that has suggested the value of different vaccine dose protocols for males and females (Fischinger et al., 2019). Females also more frequently elicit immune responses against self and are hence more likely to develop autoimmune diseases such as systemic lupus erythematosus and multiple sclerosis than males (Angum, Khan et al., 2020, Jacobsen & Klein, 2021). The varied mechanisms at play in driving these associations are not yet fully described, but differences in endocrine-immune interactions between females and males are known to contribute to sex-based differences in immune responses (Klein, 2000, Oertelt-Prigione, 2012).

Sex-based differences in the immune system have also been reported in rhesus macaques, a popular model system used to study immune responses given genetic similarity to humans. For example, one rhesus macaque study reported a lower infection rate among unvaccinated females as compared to males following challenge with Simian-Human Immunodeficiency Virus (SHIV), and vaccine efficacy that was only observed among male animals (Lu, Guerin et al., 2021, Om et al., 2020). A study designed to determine if complement lysis of Simian Immunodeficiency Virus (SIV) or SIV-infected cells represent a correlate of protection against SIV infection demonstrated that induction of antibodies capable of directing complement lysis post vaccination differed between males and females (Miller-Novak, Das et al., 2018). The study demonstrated that antibody-dependent SIV lysis correlated with reduced risk of infection in vaccinated males, particularly in gp140 immunized males, but not in females. Vaccinated males in the study had higher env-specific IgM titers than vaccinated females. A previous SIV vaccine efficacy study that focused on exploring immunogenicity and protective efficacy of monomeric SIV gp120 with oligomeric SIV gp140 in rhesus macaques demonstrated that females exhibited delayed SIV acquisition as compared to males associated with enhanced mucosal B cell responses at the site of virus exposure (Tuero, Mohanram et al., 2015). Subsequent study demonstrated that several immune parameters developed differently in vaccinated males and females, including T_FH,_ env-specific antibody levels, and B_reg_ frequencies. (Mohanram, Demberg et al., 2016, Vargas-Inchaustegui, Demers et al., 2016). Such sex-associated differences may be widespread, as a study in Chinese rhesus macaques showed higher levels of leukocyte sub-populations in females (Xia, Zhang et al., 2009).

In contrast to humoral and cell-based immunity, sex-based differences in complement pathway-mediated immunity are less studied, though some differences have been reported (Rowe, Wu et al., 2020). The complement system is an innate immune surveillance system comprised of both soluble and membrane bound proteins. The system can be activated by three distinct initiation events—classical, alternative and lectin pathways, each leading to a common terminal pathway. Recognition of pathogen-associated molecular patterns or binding of complement protein C1q to the Fc portion of IgG or IgM antibodies complexed with antigen can induce the activation of classical complement pathway. The triggering force behind activation of lectin pathway is recognition of pathogen-associated carbohydrates by mannose binding lectin (MBL), followed by induction of the complement cleavage cascade through activation of MBL-associated serine proteases. The alternative complement pathway is activated when spontaneous hydrolysis of a thioester bond in complement protein C3 reaches a critical threshold (Lachmann, Lay et al., 2018, Nilsson & Nilsson Ekdahl, 2012). All three pathways give rise to formation of C3 convertase (C4b2b in classical and lectin pathway, C3bBb in alternative pathway), which catalyzes proteolytic cleavage of complement protein C3 into C3a and C3b. C3b binds to C3 convertase to form C5 convertase which cleaves protein C5 into C5a and C5b. C5b recruits complement proteins C6, C7, C8 and C9. Polymerization of C9 with the membrane leads to formation of the membrane attack complex (MAC) which forms pores that disrupt membrane integrity and lead to lysis (Merle, Church et al., 2015). Many steps in the pathway are held in check by regulatory molecules, so that activity of complement proteins is preferentially confined to appropriate pathogenic surfaces, minimizing bystander tissue damage.

Apart from the role of the complement pathway in innate immune responses which include opsonization, lysis, and generation of inflammatory responses through soluble mediators, the pathway also has a role in adaptive immunity. Studies have demonstrated that depletion of C3 impairs humoral immune responses (Pepys, 1972). Acting as a functional bridge between innate and adaptive immune responses, the complement system can enhance B cell immunity through Complement Receptor 1 (CR1) and CR2 expressed on B lymphocytes and follicular dendritic cells (Dunkelberger & Song, 2010). Complement effectors are engaged with humoral immunity at multiple stages of B cell differentiation (Carroll, 2004, Carroll, 2008), motivating consideration of its role as an ‘instructor to humoral immune response’ (Dunkelberger & Song, 2010). Antigen and immune complexes can be transported to follicular dendritic cells and to germinal centers in a complement-dependent manner (Irvine & Read, 2020, Tokatlian, Read et al., 2019). Although the precise mechanism(s) by which complement regulates T cell immunity are not known, studies have demonstrated links between complement activation and enhanced T cell responses. Antagonism of complement receptor C5aR results in fewer antigen-specific CD8 T cells following influenza A infection (Kim, Dimitriou et al., 2004), and this same receptor mediates a synergistic effect with toll like receptor 4 (TLR4) in eliciting a stronger inflammatory response (Zhang, Kimura et al., 2007). Crosslinking of complement inhibitory proteins, namely membrane cofactor protein (MCP), decay activating factor (DAF), and complement receptor CR1 can result in direct modulation APC or T cell function (Dunkelberger & Song, 2010). Crosslinking of CR1 inhibits proliferation and reduces IL-2 production (Wagner, Ochmann et al., 2006). DAF has multifaceted role in T cell biology which includes suppression of T cell responses as well as proliferation of CD4 T cells (Capasso, Durrant et al., 2006).

Given the importance of the complement system to many aspects of immunobiology, we aimed to explore the extent of similarity and differences in complement pathway-mediated immunity between males and females and in humans and rhesus macaques. Better knowledge of sex- and species-associated levels and activities of complement cascade components have the potential to not only inform study design and interpretation, but to provide insights that might contribute to advances in clinical care.

## Results

### Complement proteins are highly conserved in Humans and Rhesus

The complement system is comprised of a series of activating and suppressing proteins, expressed in soluble form as well as an on cellular membranes that interact and catalyze transformations that can either negatively regulate or result in loss of membrane integrity (**Figure 1A**). We wished to explore the conservation of human and rhesus proteins at the level of primary structure and obtained the sequences of complement proteins from humans and rhesus from the uniprot database (UniProt, 2021) (**Table 1**). Because Complement Component 7, Factor H and Factor I proteins of rhesus were not annotated, sequences for these proteins were obtained by applying BLAST (Basic Local Alignment Search Tool) algorithm (Altschul, Gish et al., 1990) using human sequences as a query on the *Macaca mulatta* proteome. The alignment score as calculated by the clustalW algorithm (Thompson, Higgins et al., 1994) revealed that complement proteins in humans and rhesus are highly conserved. Among the complement proteins considered for sequence-based conservation analysis, the lowest sequence alignment score was observed for Factor H (88.14%), and the highest was observed for C2 (97.87%). None of the complement proteins which were considered in this analysis were identical in humans and rhesus.

**Figure 1:**
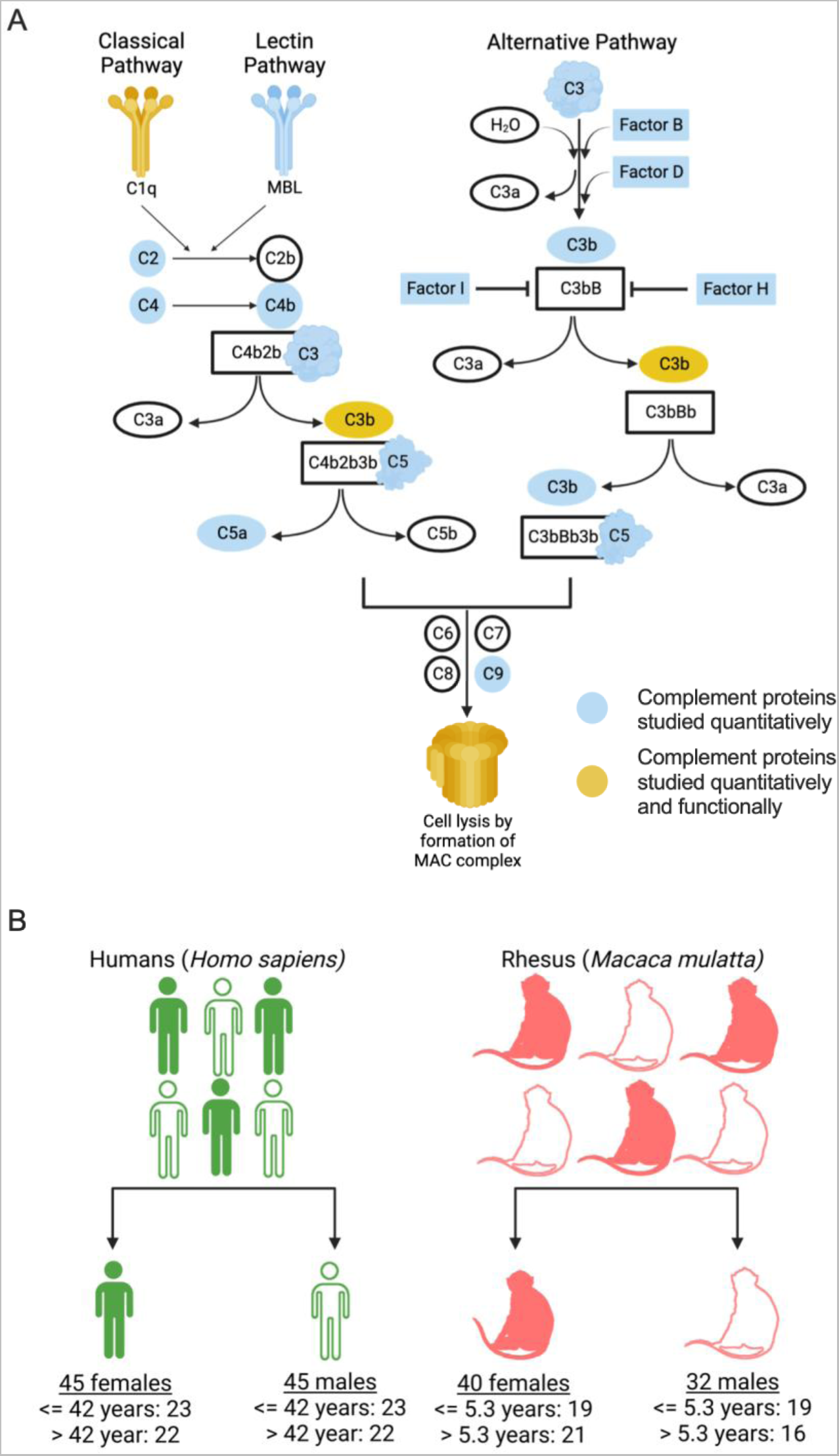
Study design. **A**. Complement Pathway with complement proteins studied quantitatively in this work are highlighted in blue and complement proteins studied quantitatively and functionally highlighted in yellow. **B**. The cohort used for the study comprised of 90 human serum samples (45 females and 45 males) and 72 rhesus serum samples (40 females and 32 males). A median split based on age was performed to categorize human and rhesus samples into two groups. Illustration created with Biorender.

**Table 1.**
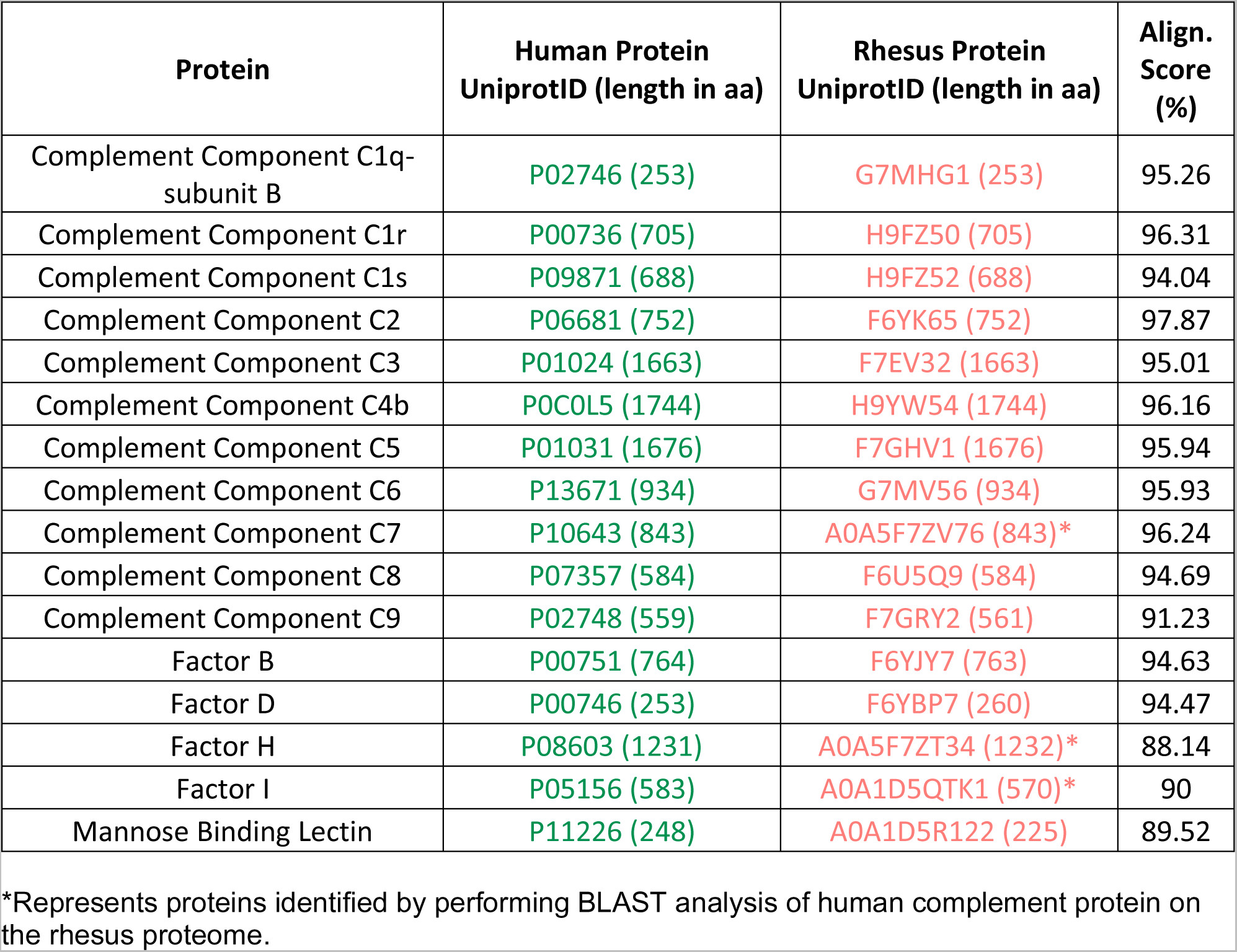
Extent of homology between Human and Rhesus complement cascade proteins studied. Pairwise global alignment of human and rhesus complement proteins using ClustalW. Alignment score was defined as the number of identities between two sequences divided by length of the alignment, and represented as a percentage.

### Complement proteins abundance in Humans and Rhesus

In humans, the levels of many of these proteins can be assessed in multiplexed assays, however, the suitability of these assays for use with non-human primate (NHP) samples has not been reported. Here, the levels of complement proteins were assessed in serum samples from all 90 human and a subset of the total 72 rhesus samples from a relatively even mix of male and female donors (**Figure 1B, Supplemental Table 1**). Higher median fluorescent signal intensity (MFI) magnitudes for all complement system analytes were observed in human serum samples as compared to rhesus (**Figure 2A**). While the high degree of amino acid sequence conservation between the human and rhesus complement proteins suggests that it is reasonable to expect the anti-human complement reagents in this experiment to cross-react with rhesus complement proteins, the many analytes that exhibited essentially undetectable signal suggests otherwise. Collectively, these results call into question the suitability of these reagents to define levels of cascade components among NHP.

**Figure 2:**
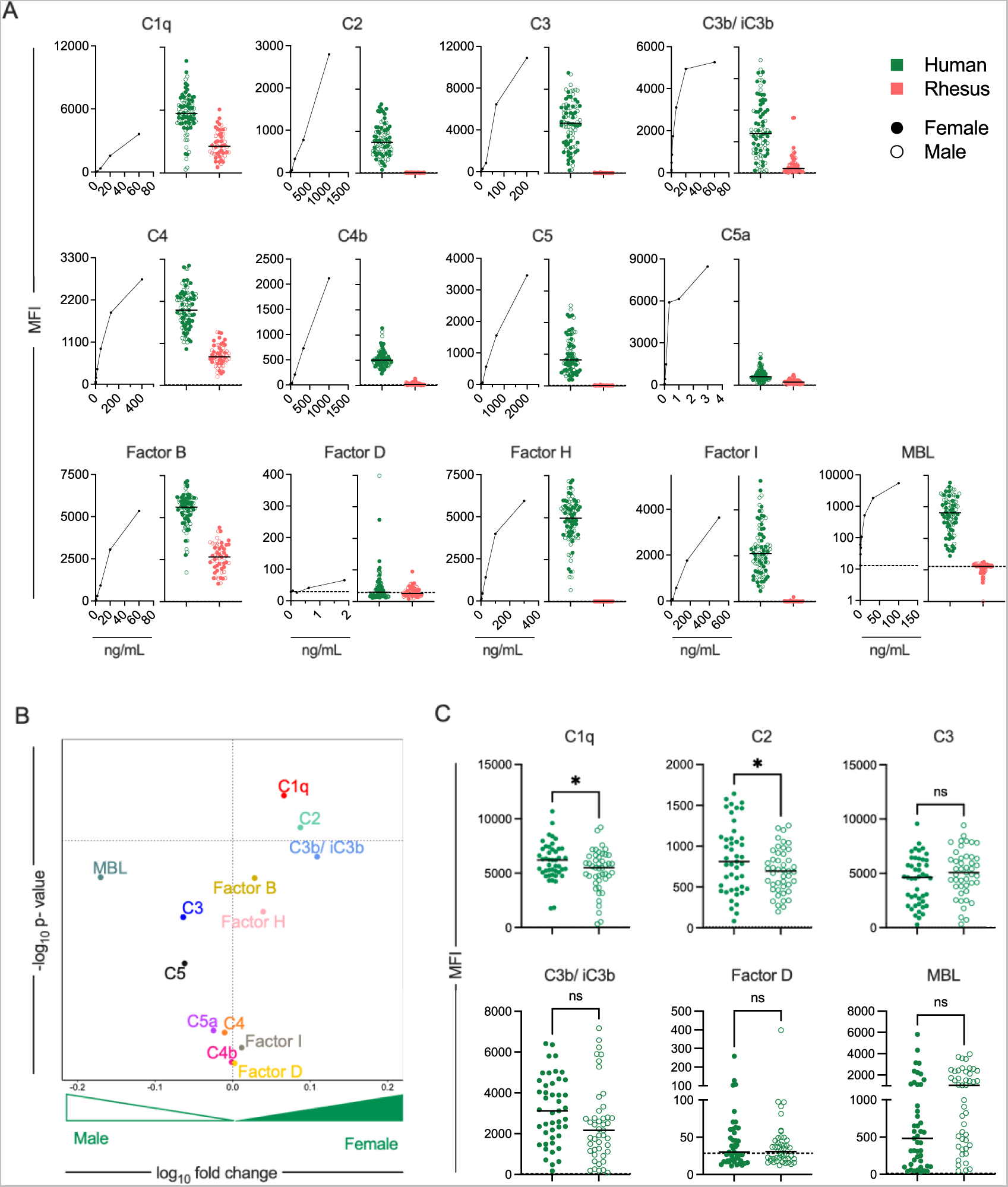
Quantification of complement proteins in Human and Rhesus serum. A. Median Fluorescent Intensities (MFI) observed in the standard curve for each human complement pathway protein (left) and for human (green) and rhesus (pink) sera samples (right) from female (solid) and male (hollow) donors. Dashed line indicates buffer only control. Bar indicates median. **B**. Volcano plot depicting the magnitude (fold-change) and statistical significance (Welch’s t-test) of differences in levels of complement proteins in human males and females. Dotted horizontal line indicates unadjusted p value of 0.05. **C**. Boxplots comparing the levels of select complement proteins in human females and males. Statistical significance defined by Welch’s t-test (*p<0.05; ns p>0.05). Dashed line indicates buffer only control. Bar indicates median.

Among humans, complement protein C1q and C2 were more abundant in female as compared to males (**Figure 2B-C**). While not statistically significant, levels of MBL and C3b were lower among females in this cohort, an observation supported by a better powered prior study among a Caucasian cohort (Gaya da Costa, Poppelaars et al., 2018). That study also reported lower levels of Factor D in females, an observation that was not supported in the primarily Hispanic and Black participant samples profiled here. Supporting these observations, analysis of B cells from individuals of age 20-30 years demonstrated overrepresentation of differentially expressed genes from the classical complement pathway in females as compared to males (Huang et al., 2021). Collectively, these results suggest the possibility of a differing balance between classical and lectin pathway activation between male and female humans.

To address age-associated differences in humans, the cohort was split at the median age (≤42 and >42 years). No age-associated differences were observed (**Supplemental Figure 1**). Previous studies among healthy humans have reported no significant differences in lectin pathway components in older (≥70 years of age) and younger adults (19-54 years of age). However, higher levels of C1q in younger (19-54 years of age) as compared to older (≥70 years of age) humans have been reported (LaFon, Thiel et al., 2021). While the current study was not designed to address differences associated with ethnicity, by chance the sample set presented an opportunity to compare the levels of complement proteins between Hispanic (n=43) and Black (n=45) donors. To this end, group differences associated with ethnicity were not observed (**Supplemental Figure 1**).

In contrast to observations in humans, sex bias in serum complement protein levels were not seen in rhesus (**Supplemental Figure 1**), though the limited signal observed for most analytes may have reduced or eliminated the ability to detect differences. While older rhesus males showed higher levels of MBL as compared to younger males (**Supplemental Figure 1**), no such difference was seen in females (**Supplemental Figure 1**).

### Human serum shows higher antibody-dependent complement deposition than Rhesus serum

Antibody-dependent deposition of complement proteins C1q and C3b was tested with two monoclonal antibodies (10-1074 (Mouquet, Scharf et al., 2012) and VRC01 (Wu, Yang et al., 2010)) specific for different epitopes on the HIV envelope (Env) glycoprotein, and pooled polyclonal serum IgGs from people living with HIV (HIVIG) using individual human and rhesus serum samples as a source of complement at two different concentrations. This assay was multiplexed across different sequence variants (n=11, **Supplemental Table 2**) of the Env glycoprotein (**Figure 3A**). In general, 10-1074 elicited higher deposition of complement proteins as compared to VRC01 (**Figure 3B**), consistent with a prior study that used more biologically-relevant lysis assays (Dufloo, Guivel-Benhassine et al., 2020).

**Figure 3:**
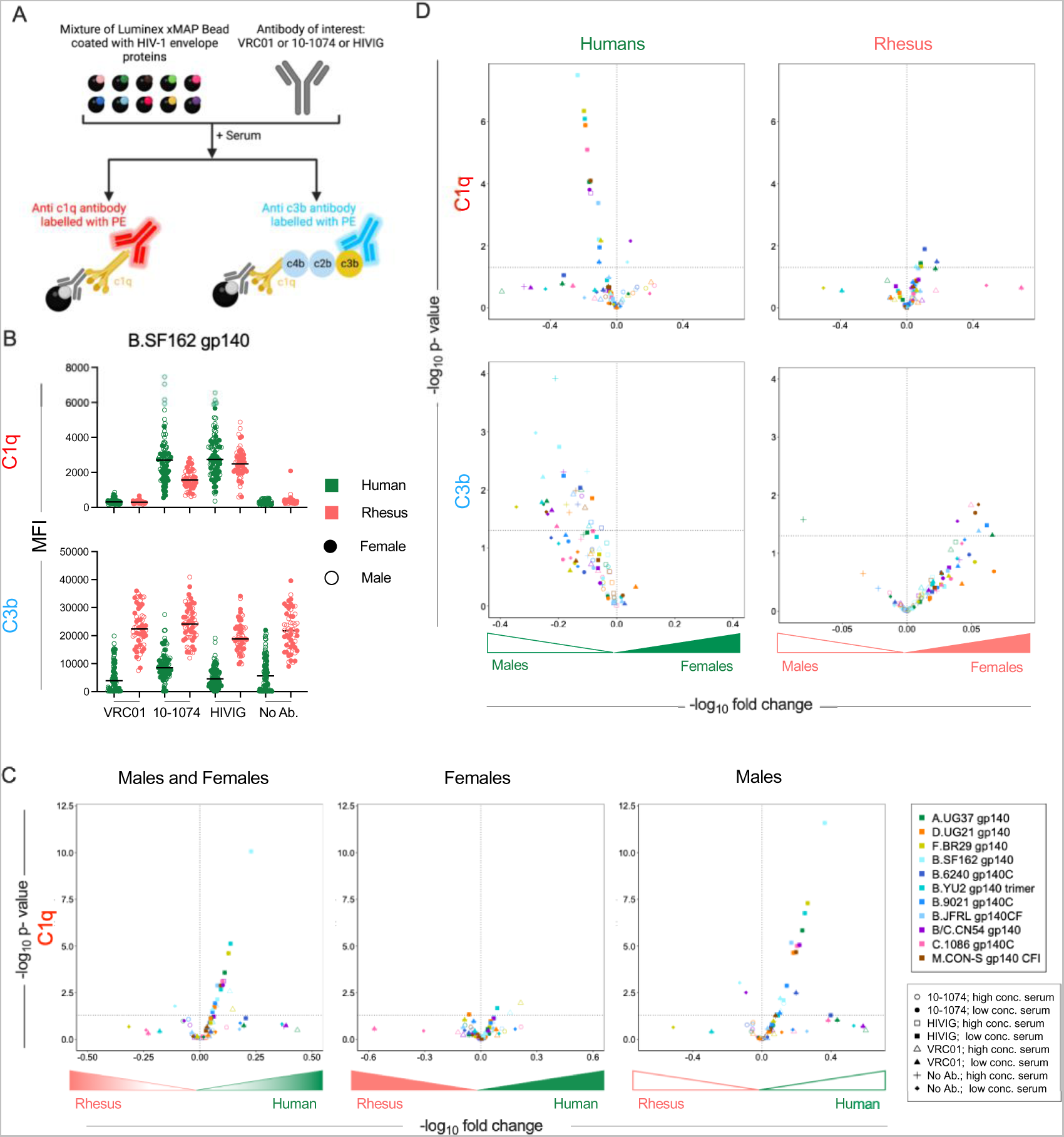
Antibody-dependent C1q and C3b deposition. A. Schematic depicting complement deposition multiplex assay in which VRC01 or 10-1074 mAbs, or polyclonal serum IgG from people living with HIV (HIVIG) were used to opsonize fluorescently-coded beads conjugated with various HIV envelope glycoproteins and assess C1q and C3b deposition following incubation with human or rhesus donor serum. Illustration created with Biorender. **B.** Box plot of total C1q (top) and C3b (bottom) detected on an envelope glycoprotein (Clade B SF162 gp140) using antibodies VRC01, 10-1074, HIVIG and no antibody control for human (green) and rhesus (pink) serum. Statistical significance defined by ordinary one-way ANOVA (****: p <= 0.0001; ns: p>0.05). **C-D.** Volcano plots of significance (Welch’s t-test) and mean fold change of antibody-dependent deposition of C1q (top) and C3b (bottom) in humans (left), and rhesus (right) for complement protein between sexes (**C**), and for C1q between species across all (left), female (center), or male (right) serum samples (**D**). Symbol shapes indicate serum concentration and antibody used and color indicates the antigen. Dotted horizontal line indicates unadjusted p=0.05.

In the absence of antibody, rhesus serum showed higher C3b deposition as compared to human serum (**Fig. 3B**). Hence, interspecies comparisons of antibody-dependent C3b deposition were not performed. Such differences were not seen in case of C1q deposition. Volcano plots were constructed to analyze differences between sexes and species. Overall, human serum showed higher antibody-dependent deposition of complement protein C1q as compared to rhesus (**Figure 3C**). Whereas C1q deposition among females was not distinct between species, elevated responses were observed for C1q in human males as compared to rhesus males (**Figure 3C**).

### Sex-based differences in complement deposition observed in Humans

We then studied sex-based differences in humans and rhesus for antibody-dependent deposition of complement proteins C1q and C3b. Despite exhibiting lower levels of C1q in serum (**Figure 1B**, **Figure 1C)**, human males showed higher deposition of complement protein C1q as opposed to human females in this assay (**Figure 3D, top left**). This degree of sexual dimorphism was not observed among rhesus (**Figure 3D, top right**). For complement protein C3b, human males showed higher deposition as compared to human females for both antibody-independent and antibody-dependent measures (**Figure 3D, bottom left**). In contrast, for a few antigens, rhesus females showed higher deposition of both C1q and C3b as compared to rhesus males (**Figure 3D, bottom right**).

No differences were seen in antibody-mediated deposition of C1q and C3b proteins in comparison of humans in association with age (**Supplemental Figure 2**). Similarly, no differences were seen in deposition of complement proteins C1q and C3b associated with race (**Supplemental Figure 2**). Age was also not generally associated with C1q or C3b deposition in rhesus; while a few measures differed for C3b deposition (**Supplemental Figure 3**). For a few antigens, higher elicitation of C3b was seen in older than younger female rhesus, while the opposite trend was observed in males (**Supplemental Figure 3**).

### Sex- and species-based differences in Antibody-dependent Complement-mediated lysis (ADCML)

Three different functional assays were performed to evaluate the ADCML activity of human and rhesus serum. In the first, lysis of sheep red blood cells (RBC) was performed to evaluate differences in ADCML elicited by human and rhesus serum by the classical complement pathway (Ross, 1986). Hemolysin was used as the source of anti-sheep erythrocyte antibody and the amount of hemoglobin released from sheep RBC post addition of serum samples was measured (**Figure 4A**). Because the surface of sheep RBC is rich in sialic acid, Factor H is able to bind to it, blocking the alternative pathway of complement activation (Chabannes, Bordereau et al., 2021). Human serum samples (n=90) demonstrated higher hemolysis of sensitized sheep RBC as compared to rhesus serum samples (n=72) (**Figure 4B**). In contrast, differences were not observed between either non-antibody-mediated hemolysis or heat-inactivated serum, which was assessed in just a few samples as a negative control, between species (**Figure 4B**).

**Figure 4.**
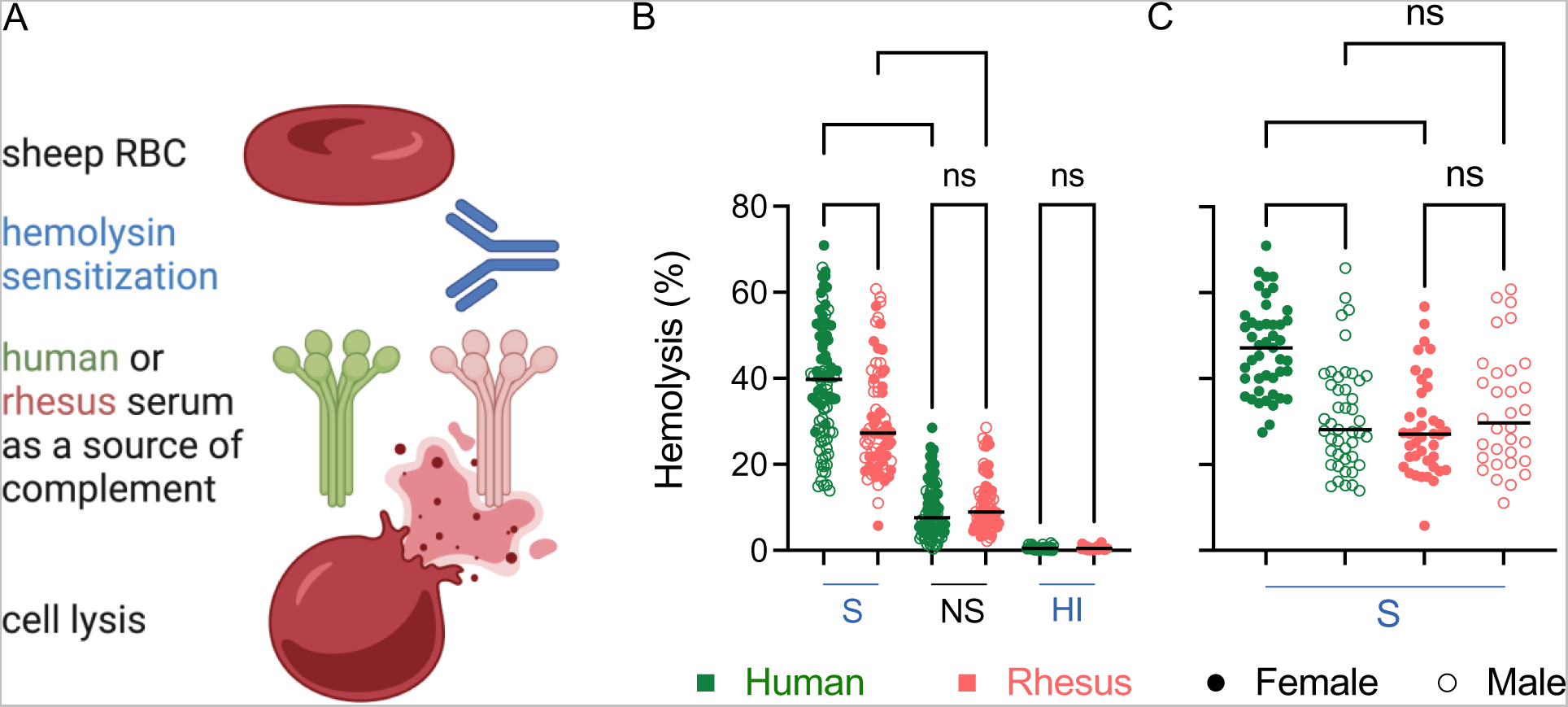
Hemolytic activity of Human and Rhesus serum. **A**. Schematic depiction of assay used to evaluate the lytic activity of human and rhesus serum against sensitized (S) sheep red blood cells (RBC). Illustration created with Biorender. **B.** Hemolysis of sheep RBCs mediated by human (green) or rhesus (pink) serum. Non-sensitized (NS) sheep RBCs and heat inactivated (HI) serum were used as controls. **C**. Sex-associated differences in hemolysis mediated by human or rhesus serum using sensitized (S) sheep RBCs. Statistical significance defined by ordinary one-way ANOVA (****p<0.0001; ns p>0.05). Bar indicates median.

The differences in hemolytic activity between species could be attributed to sex-specific differences. Whereas serum from human females (n=45) elicited greater hemolysis of sensitized RBC as compared to serum from human males (n=45), sex-associated differences were not observed in rhesus macaques (32 males, 40 females) (**Figure 4C**). Serum from human females elicited higher hemolysis as compared to serum from rhesus females, while differences between human and rhesus males were not observed (**Figure 4C**).

In both humans and rhesus, no differences in hemolysis were associated with age (**Supplemental Figure 4, Supplemental Figure 5**). Nor were differences in hemolysis associated with race in humans (**Supplemental Figure 4**). Although not statistically significant, human females >42 years of age showed higher hemolysis of RBCs as compared to human females ≤42 years of age (**Supplemental Figure 4**). Female rhesus macaques >5.3 years of age showed higher hemolysis as compared to female rhesus macaques ≤5.3 years of age (**Supplemental Figure 5**). No differences were seen in males.

### Sex- and species-based differences in Antibody-Dependent Complement-mediated lysis (ADCML) with Human and Simianized antibodies

Next, we assessed ADCML mediated by the anti-CD20 antibody Rituximab (Rtx) and its simianized equivalent (Rh Rtx), in which Rtx complementarity determining regions were grafted onto rhesus kappa light chain and IgG1 heavy chains. Antibody was added to CD20+ Ramos cells, followed by addition of serum as a source of complement. The activity of proteases released by lysed Ramos cells was measured using a luminescence-based assay (**Figure 5A**). While differences death mediated by serum between species were not observed, the Rh Rtx exhibited greater activity than Rtx, indicating greater activity of the rhesus than human IgG1 backbone, though a difference in post-translational modifications cannot be ruled out (**Figure 5B**). Consistent with the greater deposition of complement cascade factors observed in the bead assay, lysis in the absence of antibody was higher in rhesus as compared to human serum (**Figure 5B**). When the relative magnitudes of antibody-elicited cell death were compared to those observed in the absence of antibody on a per subject basis, human serum samples elicited higher ADCML as compared to rhesus serum for both antibodies (**Figure 5C**), as did serum samples from human females as compared to either human males or rhesus females (**Figure 5D**). Sex-associated differences in lysis were not observed for either antibody in rhesus macaques. Differences in activity were not observed to associate with either age in either humans or rhesus (**Supplemental Figure 4, Supplemental Figure 5**), or to associate with race in humans (**Supplemental Figure 4**).

**Figure 5.**
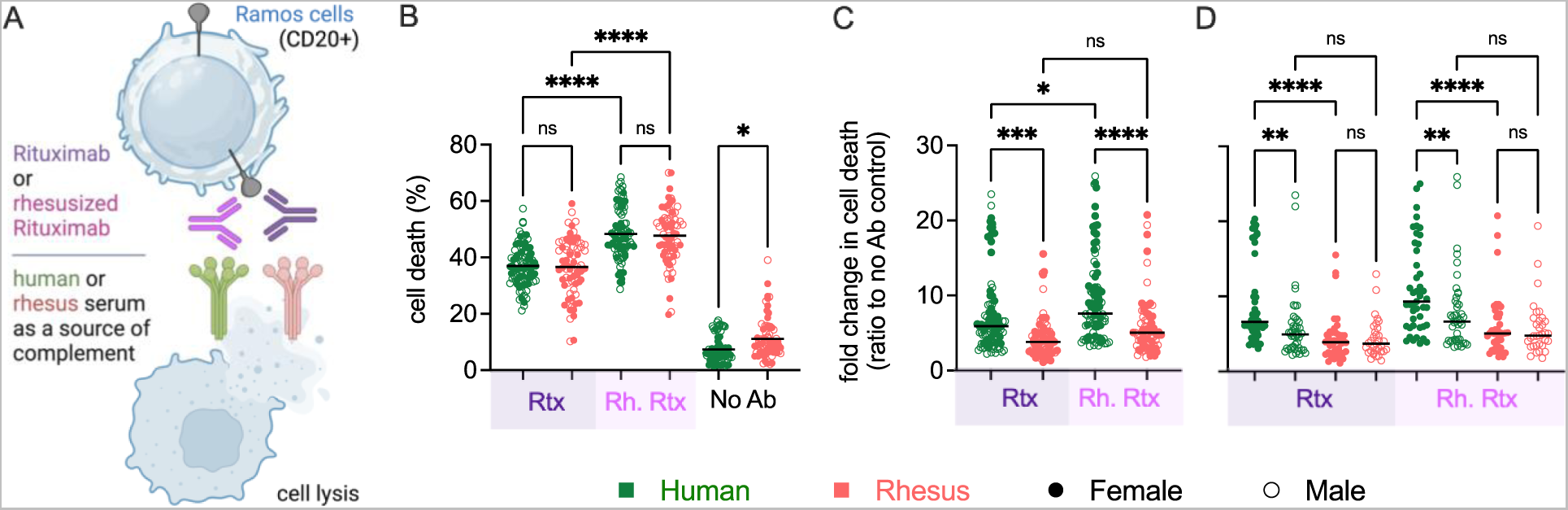
Rituximab-mediated lysis activity of Human and Rhesus serum. **A**. Schematic depiction of assay used to evaluate the lytic activity of human and rhesus serum against rituximab-treated Ramos B cells. Illustration created with Biorender. **B.** Lysis of Ramos cells using human (green) or rhesus (pink) serum in the presence and absence of human (Rtx, purple) and rhesusized (Rh. Rtx, fuschia) anti-CD20 antibody. **C**. Antibody-dependent lysis of Ramos cells. **D**. Sex-associated differences in antibody-dependent lysis. Statistical significance defined by ordinary one-way ANOVA (****p<0.0001, ***p<0.001, **p<0.01, *p≤0.05, ns p>0.05). Bar indicates median.

### Species-based differences in Antibody-dependent Complement-mediated lysis (ADCML) with Fc-engineered antibodies

Lastly, ADCML of Raji cells expressing varying surface density of HIV-1_YU-2b_ Env antigen was performed using a limited number of human (n=3) and rhesus (n=3) serum samples for multiple Fc domain engineered monoclonal antibodies (mAb) (**Figure 6A, Supplemental Figure 6**). The broadly neutralizing antibodies 10-1074 (Mouquet et al., 2012), VRC01 (Wu et al., 2010), and 10e8v4 (Kwon, Georgiev et al., 2016), which are known to exhibit complement-mediated lytic activity (Dufloo et al., 2020, Goldberg, Spencer et al., 2023, Spencer, Goldberg et al., 2022), were tested. Among the mutations in the antibodies, LALA and KA substitutions are known to reduce complement activity and EFTAE and EG have been reported to enhance complement activity (Diebolder, Beurskens et al., 2014, Goldberg & Ackerman, 2020, Hessell, Hangartner et al., 2007, Moore, Chen et al., 2010, Wirt, Rosskopf et al., 2017). Among the antibodies, 10-1074 exhibited the greatest lytic activity, VRC01 was intermediate, and 10e8v4 exhibited the lowest activity for both human and rhesus serum, and for both high and low Env expression levels (**Figure 6B**). Rhesus serum often but not always exhibited greater activity than human serum. The extent to which KA and LALA mutations reduced activity depended on mAb, antigen density, and complement source. For example, these mutations reduced the activity of 10e8v4 for both cell targets and both species, but only did so for 10-1074 in the context of low Env expression. For VRC01, greater reductions were observed for human than rhesus serum samples. EFTAE and EG mutations, which improve affinity for C1q or increase IgG hexamerization propensity, respectively, reliably exhibited improved lytic activity across mAbs, varying Env density, and for both rhesus and human serum. The extent of improvement was typically greater for mAbs and antigen density conditions for which high activity of the unmodified mAb was not observed. Collectively, consistent with other studies (Hessell et al., 2007, Hezareh, Hessell et al., 2001, Xu, Alegre et al., 2000), this data demonstrates the suitability of Fc engineering mutations designed to influence activity in humans to extend to rhesus, while at the same time reinforcing the high degree of context-dependence on complement activation.

**Figure 6.**
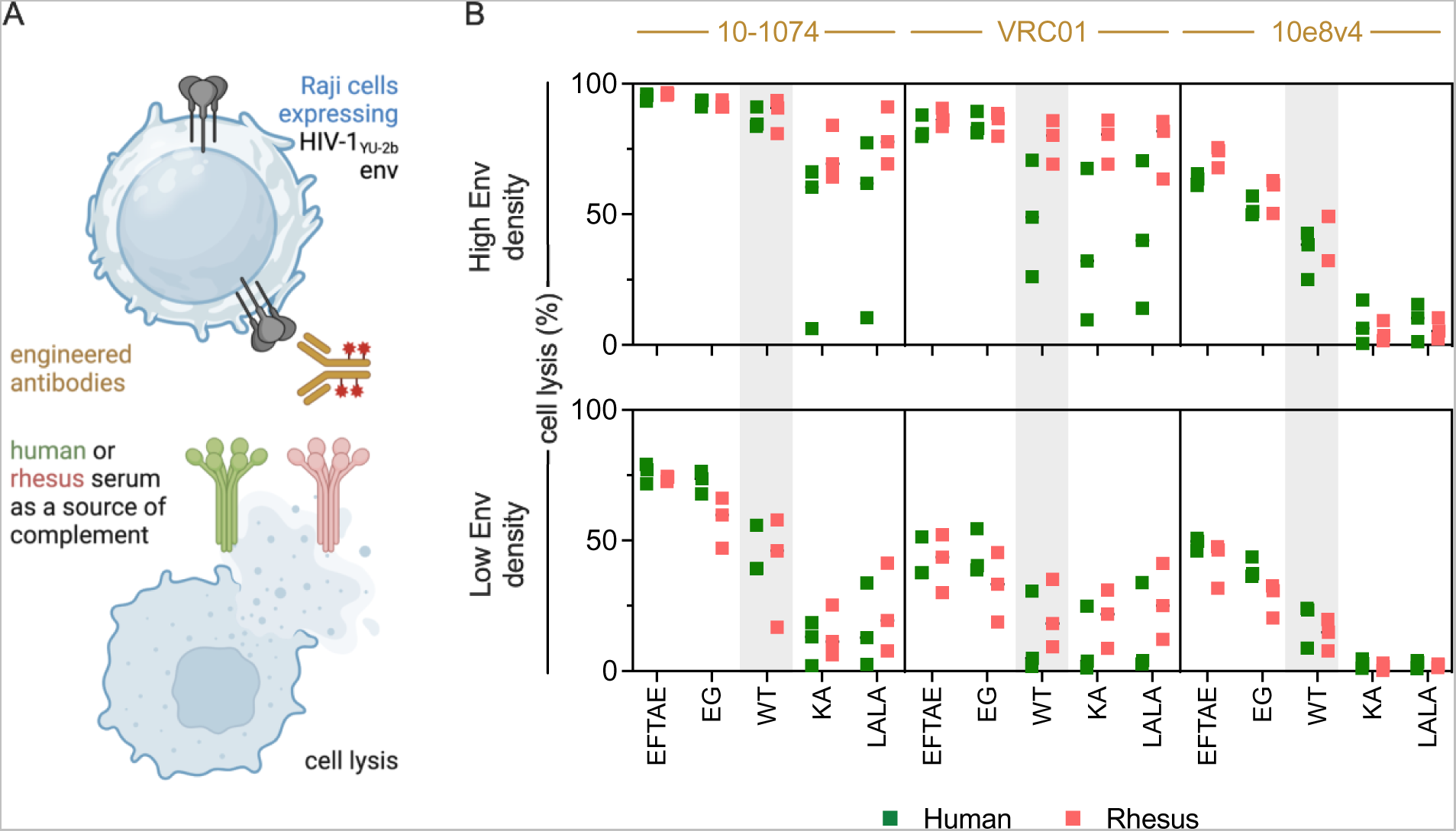
Lytic activity of Fc-engineered antibodies. **A**. Schematic depiction of assay used to evaluate HIV-1 envelope-specific lysis mediated by engineered antibodies. Illustration created with Biorender. **B**. ADCML of Raji Cells expressing high (top) and low (bottom) levels of HIV envelope glycoprotein using antibodies 10-1074 (left), VRC01 (center) and 10e8v4 (right) with and without Fc domain mutations designed to increase (EFTAE, EG) or decrease (KA, LALA) complement activation in serum from human (green, n=3) and rhesus (pink, n=3). Unmodified IgG (WT) activity is indicted in gray region in each subpanel. Percentage cell lysis was calculated with respect to a no antibody control.

### Sex- and species-based differences in Complement-aided Antibody-dependent Phagocytosis (C’ADCP) with Human and Rhesus serum

Antibody-mediated phagocytosis can be triggered by either or both Fc and Complement receptors. THP-1 cells (monocytes) express the CR3 receptor, which can recognize iC3b and drive internalization of antigenic particles (Aderem & Underhill, 1999, Tohyama & Yamamura, 2006, Vidarsson & van de Winkel, 1998). We assessed phagocytosis in the presence of complement proteins elicited by VRC01, 10-1074, as well as HIVIG (**Figure 7A, Supplemental Figure 7**). Human serum samples (n=90) demonstrated higher phagocytotic activity as compared to rhesus serum sample (n=56) for both 10-1074 and HIVIG. No difference in phagocytotic activity was seen in case of VRC01 or media alone (**Figure 7B**). Consistent with earlier observations of reduced deposition of C1q and C3b, VRC01 showed lower C’ADCP as compared to 10-1074 or HIVIG, and 10-1074 demonstrated higher C’ADCP as compared to HIVIG (**Figure 7B**).

**Figure 7.**
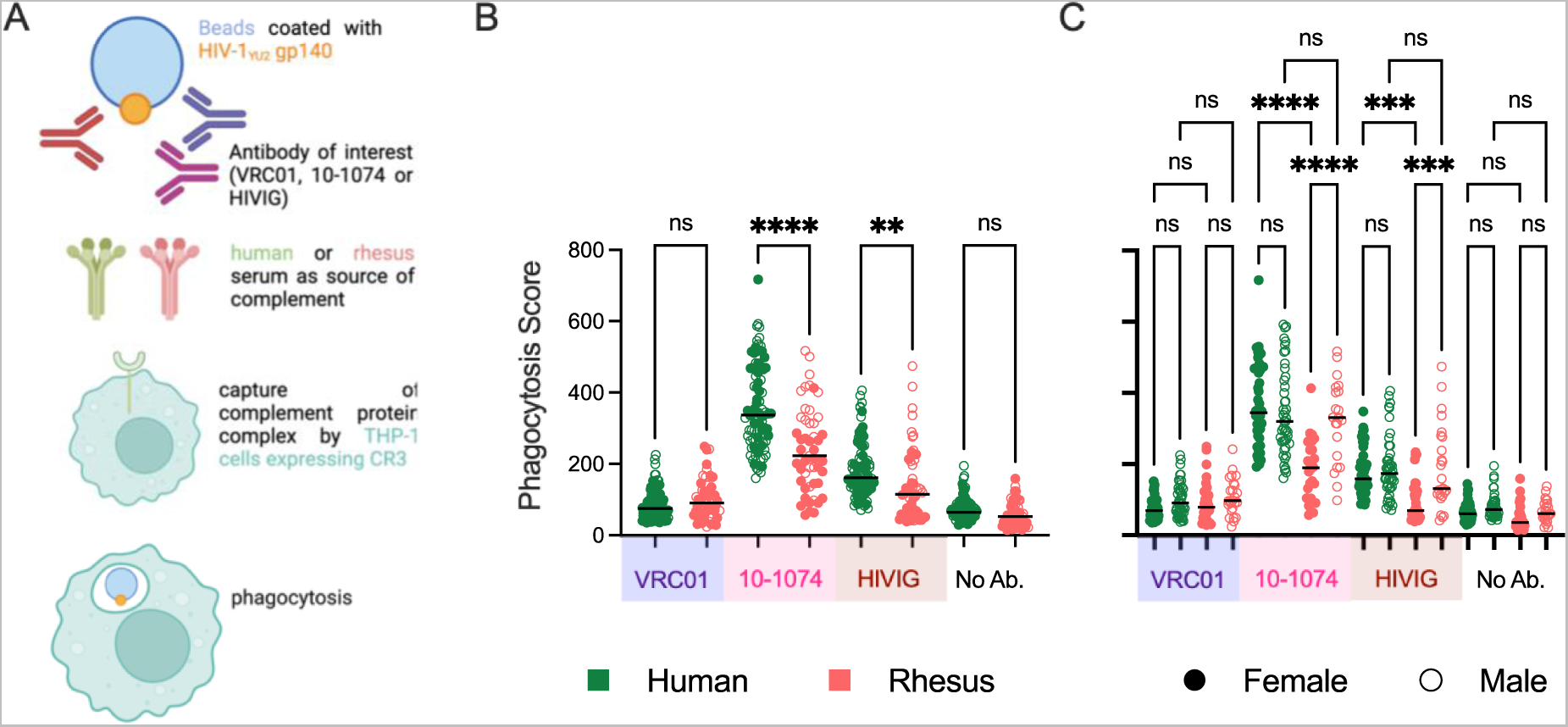
Complement-aided Antibody-dependent Phagocytotic activity of human and rhesus serum. **A**. Schematic depiction of assay used to evaluate the Complement-aided Antibody-dependent Phagocytosis (C’ADCP) by human and rhesus serum against antibodies VRC01, 10-1074 and HIVIG. Illustration created with Biorender. **B.** Box plot of phagocytosis scores of HIV-1 envelope glycoprotein (Clade B, YU2 gp140 trimer) beads using Human (green) or Rhesus (pink) serum in the presence and absence of VRC01 (mauve), 10-1074(pink), HIVIG (brown) or media alone. **C**. Sex-associated differences in C’ADCP. Statistical significance defined by ordinary one-way ANOVA (****p<0.0001, ***p<0.001, **p<0.01, *p≤0.05, ns p>0.05). Bar indicates median.

The differences in phagocytotic activity between species could be attributed to sex-specific differences. Serum from human females elicited higher phagocytotic activity than rhesus females for 10-1074 and HIVIG (**Figure 7C**). No such interspecies difference was observed in males. Whereas no difference in C’ADCP was observed among human females and human males, serum from rhesus males elicited higher C’ADCP activity as compared to serum from rhesus females for 10-1074 and HIVIG (**Figure 7C**).

Among rhesus males, in absence of antibody, males ≤5.3 years of age elicited lower phagocytotic activity as compared to males >5.3 years of age (**Supplementary** Figure 8. Age-associated differences were not seen in females. In both humans and rhesus, no differences C’ADCP were associated with age (**Supplemental Figure 8, Supplemental Figure 9**). Nor were differences in phagocytosis associated with race in humans (**Supplemental Figure 9**).

## Discussion

Sex-based differences in immune responses are known to impact the severity, pathology and prognosis of diseases. However, there is limited understanding about sex-associated differences due to complement-mediated immunity in humans and rhesus macaques, a non-human primate model organism used to study immune responses and model disease. By measuring the levels of various complement cascade proteins in serum, as well as a battery of complement activities in various functional assays, the current study sought to compare complement pathway-mediated immunity between the sexes in humans and rhesus macaques.

The resulting data suggests sexual dimorphism in complement pathway among humans as well as rhesus macaques. In humans, activation of early complement pathway components was higher in males, while antibody-mediated cell lysis was higher in females. Perhaps consistent with these observations, studies in mice have demonstrated that females were dependent on MBL-mediated initiation of complement pathway, whereas males employed C1q as well as MBL to initiate the complement cascade (Wu, Rowe et al., 2021). Such sexual dimorphism may be due to interaction of sex hormones and the complement pathway. Indeed, the promoter of the complement protein *C3* gene is responsive to stimulation by estrogen in transcriptional assays in mammalian cells(Norris, Fan et al., 1997). High levels of estrogen in females have also been shown to influence recruitment of factor H, which in turn acts as inhibitor of alternative complement pathway(Kumwenda, Cottier et al., 2022).

In females, menopause leads to secretion of low ovarian hormones like estrogen, whereas sharp decrease in testosterone levels is not seen in males(Hagg & Jylhava, 2021). Although not statistically significant, and not in perfect accord with the age of menopausal transition, older human females (>42 years of age) showed higher hemolysis of sheep RBCs as compared to younger females (≤42 years of age), a difference not seen in males. This differential observation was consistent in rhesus macaques. These results further strengthen the hypothesis of interactions between sex hormones and classical complement pathway proteins.

Although the present study demonstrates sex- and species-based differences in complement pathway-mediated immunity in humans and rhesus macaques, it has a number of limitations. The complement pathway consists of about 50 proteins, and levels of all the proteins were not measured. Although the cohorts used include a variety of subjects in different age ranges, human serum samples from individuals younger than 19 years of age or older than 77 years of age were not evaluated. Similarly, rhesus younger than 2.3 years of age or older than 21.2 years of age were not profiled. Multiplex assays developed to quantify human complement proteins were used for detection of rhesus proteins, which despite a high degree of sequence similarity, did not show detectable levels of many analytes. Samples were collected at different sites, and while care was taken to perform sample collection and handling consistently, uniformity could not be guaranteed. Genetic and environmental exposure history of human and rhesus macaque subjects was not available, and hence differences in genetic polymorphisms could not be studied. While some sex-associated differences were observed across assays, others were context-dependent, consistent with the sensitivity of complement activation to many factors. Lastly, while the current study highlights possible interaction of sex hormones and complement pathway proteins, information on hormone therapy status or hormone levels of the subjects was not available, and hence correlative relationships between sex hormones and complement protein levels or activity could not be determined.

Most of the deficiencies in complement components are inherited in an autosomal codominant pattern. Acquired deficiency defects of complement may result from increased consumption, decreased synthesis, or increased catabolism of complement proteins. Patients with complement component deficiency often present with pyogenic infections by organisms like *Streptococcus pneumoniae*, *Haemophilus influenza type B*, *Neisseria gonnorrhoeae*, *N. meningitidis*, whereas autoimmune diseases often result in secondary complement deficiencies (Wen, Atkinson et al., 2004). Plasma infusions have been occasionally used as source for deficient complement components (Steinsson, Erlendsson et al., 1989). Sex bias in complement protein levels suggests that sex matching of donor plasma may be worth considering.

As mentioned earlier, the level of complement proteins in plasma is determined by the balance of synthesis and consumption. The C1 inhibitor (C1-Inh) protein provides inherent stability to the complement pathway by inactivating C1r and C1a, leading to liberation of free C1q. Here, we observe higher basal levels of C1q in human females as compared to human males, but a higher level of antibody-dependent C1q deposition human males. These results might indicate sexual dimorphism in level of C1-Inh. C1-Inh deficiency leads to a rare disorder, hereditary angioedema, which demonstrates a greater disease burden in women as compared with men (Agostoni & Cicardi, 1992, Bork, Meng et al., 2006, Bouillet, Launay et al., 2013). Studies demonstrate that disease severity is exacerbated by increased exposure to estrogen (Banerji & Riedl, 2016), again highlighting the possible interaction of sex hormones and complement pathway proteins. Similarly, a study in participants suffering from late-stage knee osteoarthritis demonstrated that complement pathway activation may play a role in synovial vascular pathology mechanisms in males than in females. It also highlighted that females do not activate terminal complement pathway as effectively as males, regardless of the level of C5 present in synovial fluid (Sodhi, Philpott et al., 2022).

Although the complement pathway has many protective functions in immunity, imperfect regulation of complement proteins causes tissue damage in diseases like rheumatoid arthritis, age-related macular degeneration, multiple sclerosis, and ischemia reperfusion injury, among others (Carroll & Sim, 2011, Mollnes & Fosse, 1994). Failure to model or consider the impact of sex-associated differences has the potential to impact studies of these disease states, and may contribute to challenges in the “bench to bedside” extrapolation of these inhibitors to clinical applications (Neher, Weckbach et al., 2011). As an example, complement inhibitors have been tested for treatment of traumatic brain injury (Kaczorowski, Schiding et al., 1995, Leinhase, Rozanski et al., 2007, Leinhase, Schmidt et al., 2006, Sewell, Nacewicz et al., 2004). While multiple factors are certainly at play, most ischemia reperfusion studies use male mice, which exhibit differential terminal complement pathway activity from female mice in common strains (Kotimaa, Klar-Mohammad et al., 2016). Consistent with these differences, a recent study demonstrated that the complement initiation response to ischemia reperfusion injury differed in male and female mice: C1q-deficient male but not female mice were protected from injury, whereas MBL deficiency protected both male and female mice (Wu et al., 2021), suggesting a difference in the relative contributions of classical and lectin-mediated complement cascades. Similarly, Age-Related Macular degeneration is a chronic degenerative disease of the retina, caused by elevation in complement proteins. A recent study among patients with Intermediate Age-Related Macular degeneration demonstrated higher levels of complement factor B and complement factor I in females as compared to males (Marin, Poppelaars et al., 2022). These and other studies support consideration of sex-based differences in the design and interpretation of preclinical models. To this end, this study highlights sex-based differences in complement pathway mediated immunity in humans and rhesus macaques. A number of the sex-based differences observed were species specific, suggesting limitations in the ability of the macaque model to fully recapitulate human biology relevant to disease. Other differences were highly context-dependent, suggesting the continued value of functional evaluation of complement activities.

## Materials and methods

### Cell lines

Ramos and Raji cells were purchased from ATCC and maintained in RPMI-1640 containing 10% fetal bovine serum (FBS), at 37°C with 5% CO_2_. Raji cells expressing HIV-1 Envelope were generated as described previously (Dufloo et al., 2020). The cells were maintained in RPMI-1640 containing 10% FBS and 1 µg/mL puromycin (InvivoGen, ant-pr-1). The THP-1 human monocytic cell line was purchased from ATCC and maintained in RPMI-1640 supplemented with 10% FBS and 55 µM beta-mercaptoethanol at 37°C with 5% CO_2_.

### Serum sample acquisition

Human serum samples screened for HIV-1, HIV-2, Hepatitis B, Syphilis, West Nile virus, HCV were purchased commercially (BioIVT, HUMANSRMMN5 and HUMANSRMFN5). Among the samples, 50% were from females (n=45) and remaining 50% were from males (n=45). Serum samples from Indian-origin rhesus macaques were acquired from Oregon Health and Science University. Among the set of 72 samples, 55.56% (n=40) were from female and the remaining 44.44% (n=32) were from male macaques. Serum samples were collected by a method that preserves complement functional integrity. Briefly, blood was allowed to clot at 4°C and centrifuged for 10 min at 1200xg. The serum (top, clear) layer was then collected, aliquoted, and stored at -80°C within 2 hours of blood draw. Serum samples were aliquoted and stored at -80 °C to avoid repeated freeze thaw cycles.

### Sequence alignment

Amino acid sequences of complement cascade proteins were obtained from the uniprot database (www.uniprot.org). Rhesus factor H and factor I sequences were not available in uniprot database, and were instead putatively identified by homology to their human homologs by performing a BLAST (https://www.uniprot.org/blast) search on the rhesus proteome.

### Measuring levels of complement proteins in Human and Rhesus serum

Multiplex assay panels (Millipore Sigma, HCMP1MAG-19K and HCMP2MAG-19K) were used for detection of complement proteins essentially according to the manufacturer’s instructions. Serum samples were diluted 1:200 and 1:40,000 for HCMP1MAG-19K and HCMP2MAG-19K, respectively, using assay buffer as the diluent. The assay was performed by modifying manufacturer’s instruction to perform assay in 384 well plate as published by Tang *et al*. (Tang, Panemangalore et al., 2016). Plates (Greiner bio-one, 781906) were blocked with 50 μL of PBS containing 1% BSA for 10 min at room temperature with shaking (1000 rpm). Serum samples were diluted in in cold Gelatin Veronal Buffer containing Ca^2+^ and Mg^2+^ (GVB++) (Complement Technology, B100) and were then added to the 384 well plate. As per the manufacturer’s instruction, 6 μL of premixed beads conjugated with antibodies specific to complement proteins were added to each well containing the serum samples. The plate was sealed, wrapped with foil and was incubated with agitation at 1000 rpm on a plate shaker for 2 hours at room temperature. The plate was washed using a plate washer according to manufacturers’ instructions and 6 µL of provided detection antibody was added. The plate was sealed, covered with foil and was incubated with agitation for 1 hour at room temperature, followed by the addition of 6 µL of Streptavidin-Phycoerythrin. The plate was again sealed, covered with a foil and was allowed to incubate with agitation at 1000 rpm on plate shaker for 30 min at room temperature prior to washing and resuspension with 50 µL of sheath fluid (Luminex™ xMAP Sheath Fluid Plus, 4050021), sealed (Eppendorf, Catalogue no. 0030127854), and agitated at 1000 rpm for 5 minutes. Data was acquired on a Luminex™ FLEXMAP 3D™ Instrument System, which detected the beads and measured PE fluorescence in order to calculate the median fluorescent intensity (MFI) level for each analyte. Heat-inactivated pooled human sera (Sigma-Aldrich, S1764) and rhesus sera were used as negative controls. Heat inactivation was performed by heating serum at 58°C for 30 min. The standards in the kits were used as positive controls.

### Antibody-dependent deposition of Complement proteins C1q and C3b

Recombinant extracellular domains of HIV envelope proteins [NIH HIV Reagent Program, **Supplemental Table 2**] were covalently coupled to Luminex™ Magplex® magnetic microspheres using two step carbodiimide chemistry as described previously (Brown, Dowell et al., 2017). 384 well plate was blocked using PBS + 1% BSA as described earlier, followed by a GVB++ wash (Complement Technology, B100). HIVIG [NIH HIV Reagent Program, ARP-3957], bnAb 10-1074 IgG1 [heavy and light chains were cloned in pCMV vector, and the antibody was expressed in HEK Expi cells (Gibco™, A14635) following manufacturer’s instruction], bnAb VRC01 IgG1 [heavy and light chain plasmids for antibody expression (NIH AIDS Reagent Program, ARP-12035 and ARP-12036) were used to express the antibody in HEK Expi cells (Gibco™, A14635)], or buffer alone were added to microspheres coupled with each HIV-envelope protein were diluted in assay buffer to 60 beads/well of each antigen, and incubated with shaking for 2 hours at room temperature. All the dilutions were performed in ice cold GVB++. Following antibody binding, the plate was washed on a plate washer using PBS + 0.1% Tween20 and the complement deposition activity of sera at four final dilutions, 1:40, 1:400, 1:4000, and 1:40,000, was assayed following incubation at 37°C for 30 min. The plate was washed using PBS plus 0.1% Tween20 using an automatic plate washer. For detection of C1q, 37 µL of 1:100 dilution of biotin-anti-human-C1q (Quidel, A700) was added. The antibody and beads were mixed by sonication and vortexing to reduce bead clumping. The plate was sealed and wrapped and then allowed to incubate for 1 hour at room temperature with agitation at 1000 rpm. Post washing, 37 µL of 1:500 SA-PE (Agilent Technologies, PJ31S-1) was added as a detection agent. The plate was sonicated and vortexed and then allowed to incubate at room temperature, 1000 rpm for 1 hour. For C3b detection, 1:300 dilution of ms-anti-human-C3b (Cedarlane Labs, CL7636AP) was added as primary antibody while goat anti-ms-IgG-PE, human adsorbed (Southern Biotech, 1010-09) was added as detection reagent. The plate was washed as using PBS + 0.1% Tween20 using an automatic plate washer. The beads were suspended in 50 µL of sheath fluid and data acquired as described above.

### Sheep RBC Hemolysis Assay

The RBC hemolysis assay protocol was adapted from protocol published by Costabile *et al*. (Costabile, 2010). A volume of 1.0 mL GVB++ (Complement Technology, B100) was added to 0.5 mL 100% packed sheep RBCs (Innovative Research, ISHRBC100P15ML) and mixed by inversion, followed by centrifugation at 600g for 5 min. Supernatant was discarded and this process was repeated two more times. After the final wash, cells were centrifuged at 900g for 5 min to pack them, decanted, and resuspended in 4.5mL GVB++ to make a 10% solution of sheep RBCs. To this solution, an equal amount of 1:50 diluted Hemolysin (Complement Technology, Hemolysin) was added drop wise while swirling the tube. The tube was incubated at 30°C for 30 min with a brief swirl at 15 min. Hemolysin-sensitized RBCs were stored at 4°C till the assay was performed. The assay was performed within 24 hours of sensitizing the RBCs.

A volume of 50 µL of treated sheep RBCs were added to a V bottom 96 well plate (USA Scientific, 18339600). To this, 50 µL of 1:100 diluted serum was added. As a control to define total lysis, 50 µL of distilled water was mixed with 50 µL treated RBC, and as a no lysis control, 50 µL GVB++ was added to 50 µL treated sheep RBCs. Heat-inactivated serum samples from pooled human and rhesus serum were used as additional negative controls. Heat inactivation was performed by heating serum at 58°C for 30 min. The plate was incubated in 37°C, 5% CO_2_ incubator for 30 min. Following centrifugation at 1500g for 5 min, 50 µL from each well was transferred into a 96 well white clear bottom polystyrene plate (Costar®, 3610) and diluted with 50 µL of water. Absorbance was measured at 540 nm using SpectraMax Paradigm Plate reader (Molecular Devices). Mean lysis across triplicates was calculated using the following formula:

Percentage lysis= [OD_540_(test)-OD_540_(blank)]/[OD_540_(total lysis)-OD_540_(blank)] x 100. Data was collected for biological triplicates, and the average percentage lysis reported.

### Rituximab-induced Complement-mediated Lysis of Ramos cells

Ramos cells were washed with GVB++(Complement Technology, B100) and seeded at 1 X 10^5^ cells per well in a 96 well white clear bottom polystyrene plate (Costar®, 3610), to which 50 µL of 10 µg/mL of Rituximab (AbMole Bioscience, M5219), rhesusized rituximab (NHPRR, 2B8R1), or GVB++ was added. The plate was subjected to orbital shaking at 200 rpm for 2 min. Following antibody binding, 50 µL of a 1:100 dilution of serum was added. Following repeating the shaking procedure, the plate was incubated 37°C in a 5% CO_2_ incubator for 1 hour, iced for 10 min, and then 50 µL of CytoTox-Glo™ Cytotoxicity assay reagent (Promega, G9291) was added. The plate was shaken at 900 rpm for 1 min on an orbital shaker and then incubated for 15 min at room temperature. Luminescence dependent on serum complement was measured using SpectraMax Paradigm Plate reader, and total luminescence was measured following addition of 50 µL of lysis reagent and incubation at room temperature for 15 min. Relative Antibody-dependent lytic activity was calculated as the ratio of luminescence of antibody dependent complement-mediated cell death to luminescence observed for the no antibody control condition. Mean values of assay duplicates were reported.

### Effect of engineered antibodies on Antibody-dependent Complement-mediated Lysis (ADCML)

Antibody-dependent Complement-mediated lysis of Raji cells transduced with a retroviral vector encoding HIV-1_YU-2b_ Env was assessed as described previously (Dufloo et al., 2020). ADCML was assessed for bnAbs 10-1074 IgG1 (heavy and light chains were cloned in pCMV vector), 10e8v4 IgG1 (heavy and light chains were cloned in pCMV vector) and VRC01 IgG1 (heavy and light chain plasmids for antibody expression (NIH AIDS Reagent Program, ARP-12035 and ARP-12036)). Mutations in the wild type heavy chain plasmid were performed using site directed mutagenesis (Agilent, 200524) following the manufacturer’s protocol to generate EG, KA and EFTAE mutants (Diebolder et al., 2014, Goldberg & Ackerman, 2020, Hessell et al., 2007, Moore et al., 2010, Wirt et al., 2017). Antibodies were expressed in HEK Expi cells (Gibco™, A14635) following manufacturer’s instruction.

For the ADCML assay, Raji cells expressing variable levels of surface Env were used (D2 clone and D4 clone)(Dufloo et al., 2020). Cells were mixed with 50% serum and 15 µg/mL IgG and incubated for 24 hours at 37°C. Complement-mediated lysis was measured by staining with live/ dead fixable aqua dead cell marker (Life Technologies, Catalogue no. L34957) prior to fixation. Flow cytometric analysis (Invitrogen, Attune NxT) of the fixed and stained cells was performed. CDC was calculated using the following formula:

Percentage lysis= [% of dead cells with antibody - % of dead cells without antibody]/[100 - % of dead cells without antibody] X 100. Negative values were set to zero.

### Complement-aided Antibody-dependent Phagocytosis (C’ADCP)

To gain insight into sex/species differences in complement activity, the aggregate opsonophagocytic activity of antibody-coated, complement-fixed particles was assessed by performing an antibody-dependent cellular phagocytosis assay (Ackerman, Moldt et al., 2011) modified to include incubation with complement active serum. C’ADCP was tested with VRC01, 10-1074, and HIVIG antibodies, as described above. Antigen beads were prepared by conjugating 50 µg HIV envelope protein gp140 SF162 (NIH HIV Reagent Program, ARP-12026) was to a 400 µL volume of fluorescent beads (1.7-2.2-micron SPHERO Carboxyl Fluorescent Particles, Spherotech Inc, CFP-2052-2) using carbodiimide chemistry as described previously (Chu, Crowley et al., 2020). Serum samples were rapidly thawed, placed on ice, and subsequently distributed to 384 well plates (Greiner bio-one, 781906). The serum samples were diluted to 1:72 in GVB++ and 50 µL of diluted serum was added to 96 well V bottom plate (Plateone®, 1833-9600).

Microspheres (1x10^8^) were incubated with 100 nM of antibody (diluted in GVB++ (Complement Technology, B100)) for 2 hours at 4°C with end-over-end mixing. Following the incubation, the microspheres were washed once, resuspended, added to the V-bottom plates on ice (final serum concentration= 1:144). Plates were incubated at 37°C (orbital shaking, 800RPM) for 30 min, then placed on ice to stop the reaction before centrifugation at 4000g for 5 min. After decanting, beads were resuspended in 100 µL of RPMI 1640 + 10% FBS, and finally transferred to flat bottom 96 well tissue-culture plates (Costar®, 09-761-145), to which 100 µL of THP-1 cells (viability > 90%) at a concentration of 1 x10^5^ cells/mL were added to each well, resulting in a final effector (THP-1 cells) to target (bead coated with antibody and fixed with complement) ratio was 1:5. The effectors and target were allowed to incubated at 37°C overnight prior to fixation with 4% paraformaldehyde. The number of beads phagocytosed were measured by flow cytometer on a MACSQuant (Miltenyi Biotec) instrurment. Phagocytosis score was derived as an integrated MFI by multiplying the percentage of fluorescent (bead positive) THP-1 cells by the median fluorescent intensity of positive cells.

## Data analysis

Data was analyzed and graphed using Graph Pad Prism (Version 9.4.1), Rstudio (version 4.2.1), FloJo (BD Biosciences). Packages ggplot2 (Wickham, 2016), tidyverse (Wickham, 2019), dpylr (Wickham, 2023), ggpubr (Kassambara, 2023) were used to analyze data and generate some of the graphs.

## Acknowledgements

HIVIG was obtained through NIH HIV Reagent Program, Division of AIDS, NIAID, NIH: Polyclonal Anti-Human Immunodeficiency Virus Immune Globulin, Pooled Inactivated Human Sera, ARP-3957, contributed by NABI and National Heart Lung and Blood Institute (Dr. Luiz Barbosa). VRC01 heavy and light chain in expression vector plasmids (ARP-12035,12036) were also obtained through NIH HIV reagents program: contributed by Josh Mascola. Some of the HIV-1 gp140 antigens produced in CHO cells lines (ARP-12063, 12064, 12065, 12066) were obtained from NIH HIV reagent program: contributed by DAIDS/NIAID; produced by Polymun Scientific Inc. A few other HIV-1 gp140 antigens produced in HEK 293F cells (ARP-12572, 12573, 12575, 12577, 12581) were obtained from NIH HIV reagent program: contributed by Dr. Barton F. Haynes and Hua-Xin Liao. HIV-1 gp140 protein for SF162 envelope ectodomain produced in HEK293T cells was obtained from NIH HIV reagent program: contributed by Dr. Leo Stamatatos. Plasmid for production of HIV gp140 protein trimer from strain YU2 (ARP-12133) was obtained from NIH HIV reagent program: contributed by Dr. Joseph Sodroski. Thesimianized anti-CD20 (Rh. Rtx) (2B8R1) was obtained from NIH Non-human Primate Reagent Resource (NHPRR).

## Author contributions

N.S.K, B.S.G, J.D., T.B. collected the experimental data. A.J.H. selected and provided the rhesus macaque serum samples. O.S. and M.E.A. supervised the research. N.S.K. analyzed the data, prepared figures and drafted the manuscript. N.S.K and M.E.A finalized the manuscript. All authors reviewed and edited the manuscript.

