## Supplemental Materials for "Sex and species associated differences in Complement-mediated immunity in Humans and Rhesus macaques"

| Supplemental Figures and Legends | |
|---|---|
| Supplemental Figure 1 | Comparison of levels of complement proteins in Human and Rhesus sera by sex and age. |
| Supplemental Figure 2 | Comparison of antibody-dependent deposition of complement proteins C1q and C3b by Human serum between study subgroups. |
| Supplemental Figure 3 | Comparison of antibody-dependent deposition of complement proteins C1q and C3b by Rhesus serum between study subgroups. |
| Supplemental Figure 4 | Box plots depicting Antibody-dependent Complement-mediated Lysis (ADCML) by Human serum. |
| Supplemental Figure 5 | Box plots depicting Antibody-dependent Complement-mediated Lysis (ADCML) by Rhesus serum. |
| Supplemental Figure 6 | Gating strategy for lytic activity of Fc-engineered antibodies. |
| Supplemental Figure 7 | Gating strategy for Complement-aided Antibody-dependent Phagocytosis (C'ADCP). |
| Supplemental Figure 8 | Box plots depicting Complement-aided Antibody-dependent Phagocytosis (C'ADCP) by Rhesus serum. |
| Supplemental Figure 9 | Box plots depicting Complement-aided Antibody-dependent Phagocytosis (C'ADCP) by Human serum. |
| **Supplemental Tables and Legends** | |
| Supplemental Table 1 | Human and Rhesus Serum sample cohort |
| Supplemental Table 2 | Antigens used in multiplex assay |
| **Supplemental Data Files** | |
| Data File 1 | Level of Complement proteins in human and rhesus sera |
| Data File 2 | Antibody-dependent deposition of complement protein C1q by Human and Rhesus serum |
| Data File 3 | Antibody-dependent deposition of complement protein C3b by Human and Rhesus serum |

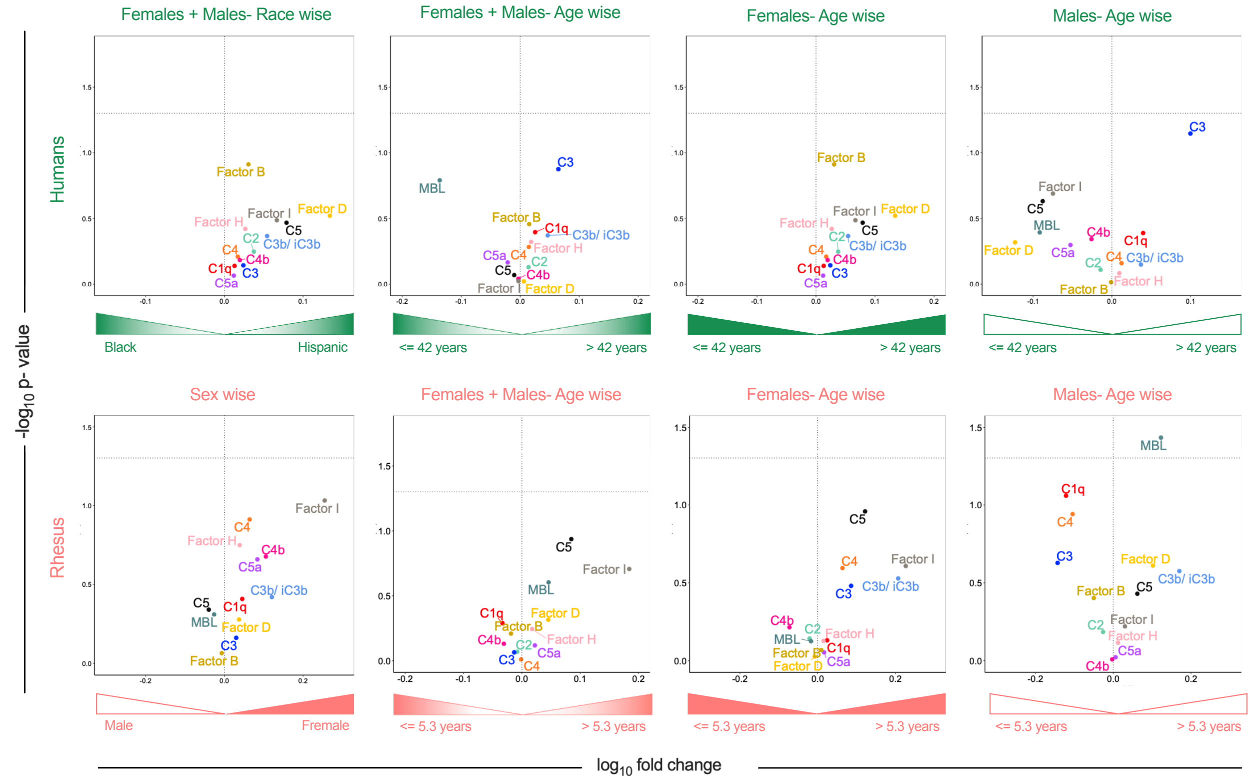


**Supplemental Figure 1: Comparison of levels of complement proteins in Human and Rhesus sera by sex and age.** Volcano plots of significance (Welch's t-test) and mean fold change of complement protein levels for indicated comparisons amongst humans (top) and rhesus (bottom). Dotted horizontal line indicates unadjusted p=0.05.

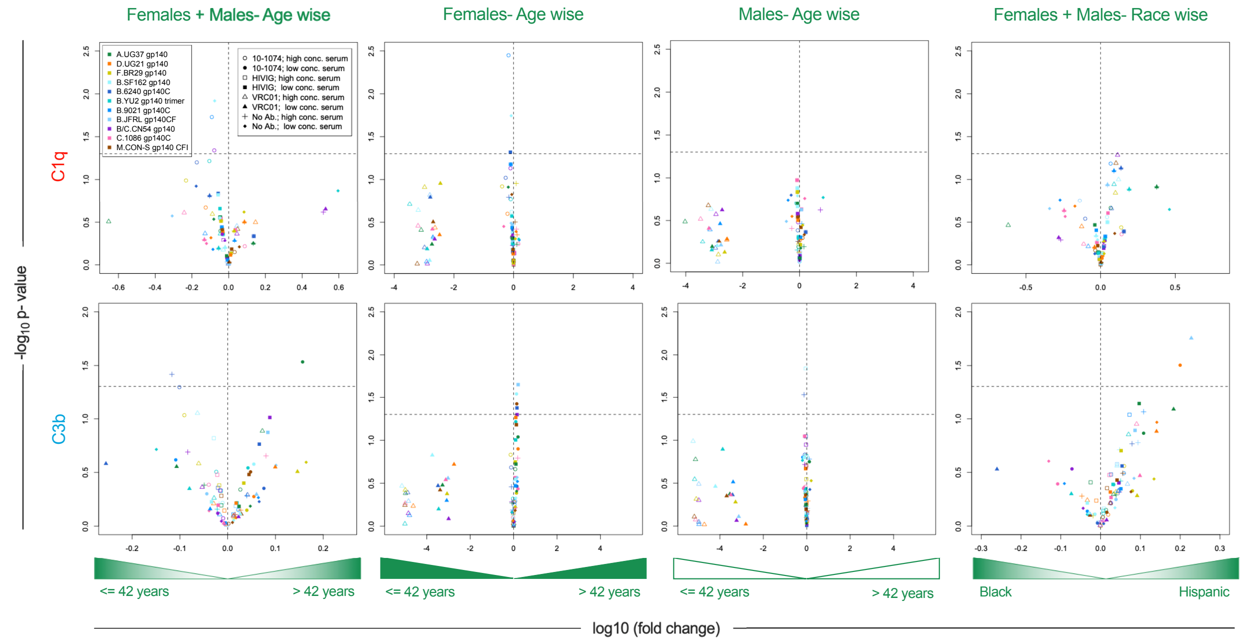


**Supplemental Figure 2: Comparison of antibody-dependent deposition of complement proteins C1q and C3b by Human serum between study subgroups.** Volcano plots of significance (Welch's t-test) and fold change of antibody-dependent deposition of C1q (top) and C3b (bottom) across HIV envelope proteins between indicated subgroups. Symbol shapes indicate serum concentration and antibody used and color indicates antigen. Dotted horizontal line indicates unadjusted p=0.05.

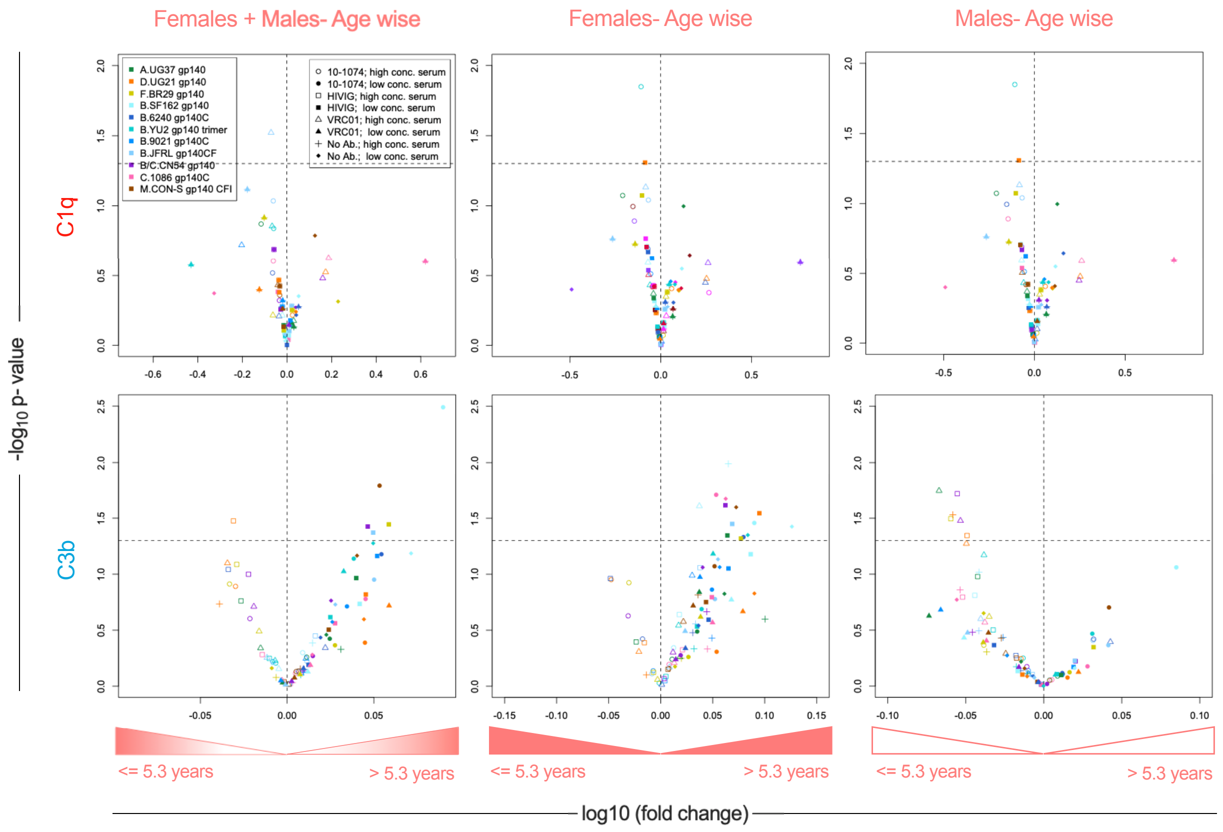


**Supplemental Figure 3: Comparison of antibody-dependent deposition of complement proteins C1q and C3b by Rhesus serum between study subgroups.** Volcano plots of significance (Welch's t-test) and fold change of antibody-dependent deposition of C1q (top) and C3b (bottom) on HIV envelope antigens between indicated subgroups. Symbol shapes indicate serum concentration and antibody used and color indicates antigen. Dotted horizontal line indicates unadjusted p=0.05.

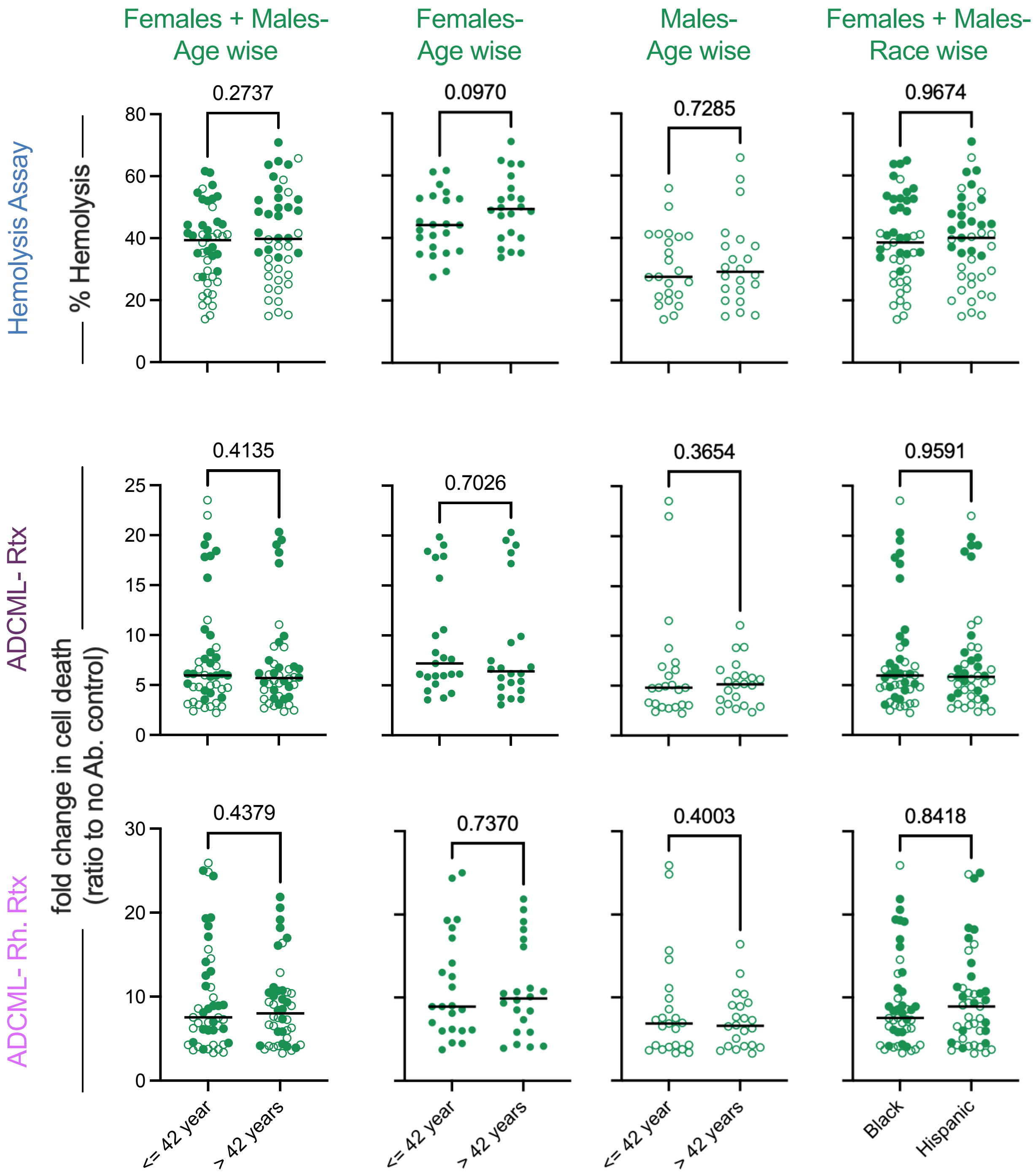


**Supplemental Figure 4: Box plots depicting Antibody-dependent Complement-mediated Lysis (ADCML) by Human serum.** Box plots of ADCML elicited by human serum based on age of Females+ Males, age of Females, age of Males and race wise as measured by Sheep RBC Hemolysis Assay (top), cytotoxicity elicited by Rtx. (center) and cytotoxicity elicited by Rh. Rtx (bottom). The percentage cell death was normalized to cell lysis in case of no antibody for Rtx and Rh. Rtx. Statistical significance defined by unpaired t test using Welch's correction. Bar indicates median.

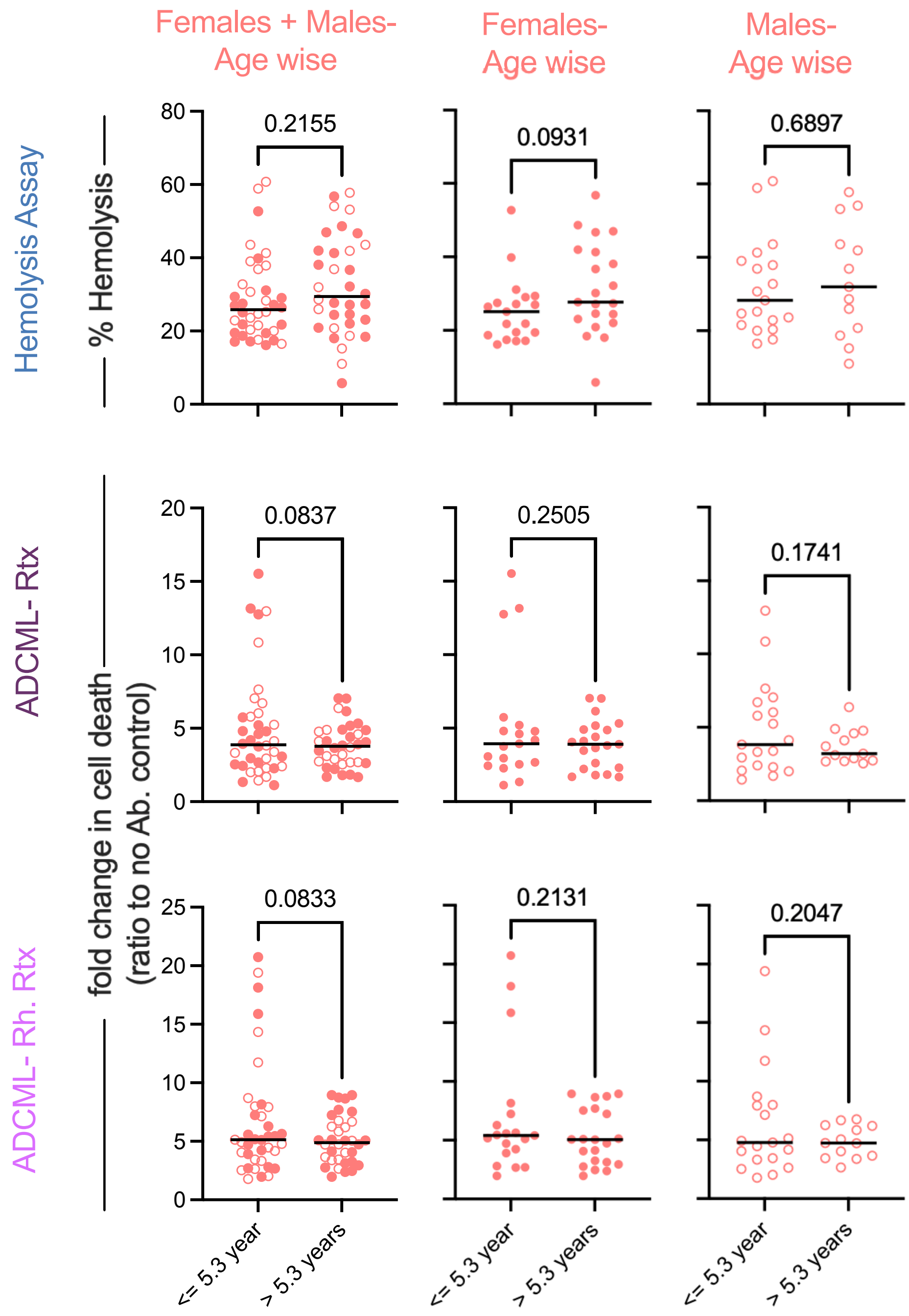


**Supplemental Figure 5: Box plots depicting Antibody-dependent Complement-mediated Lysis (ADCML) by Rhesus serum.** Box plots depicting ADCML elicited by rhesus serum based on age of Females + Males (left), age of Females (center) and age of Males (right) as measured by Sheep RBC Hemolysis Assay (top), cytotoxicity elicited by Rtx. (center) and cytotoxicity elicited by Rh. Rtx. (bottom). The percentage cell death was normalized to cell lysis in case of no antibody for Rtx. and Rh. Rtx. Statistical significance defined by unpaired t test using Welch's correction. Bar indicates median.

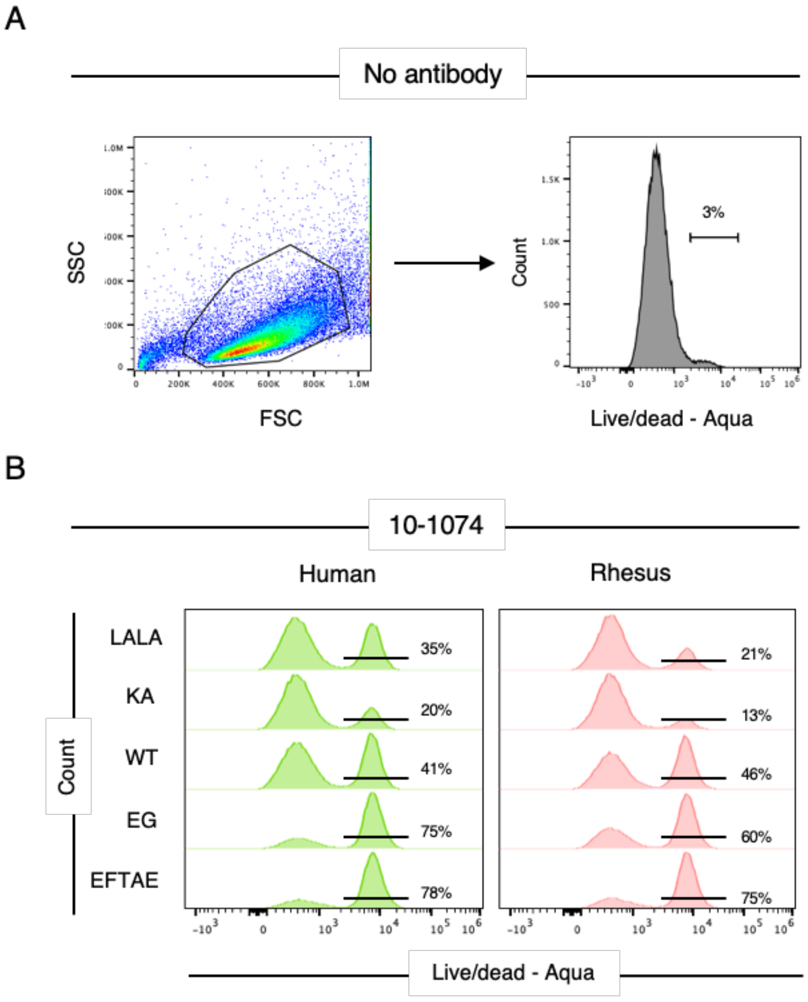


**Supplemental Figure 6: Gating strategy for lytic activity of Fc-engineered antibodies. A**. The FSC/SSC gate was drawn to include the Raji cells. The gate for live dead aqua stain was set using the information for no antibody control. **B**. Histograms for live dead staining for one of the human serum sample, and one of the rhesus serum sample for 10-1074 wild type and mutants.

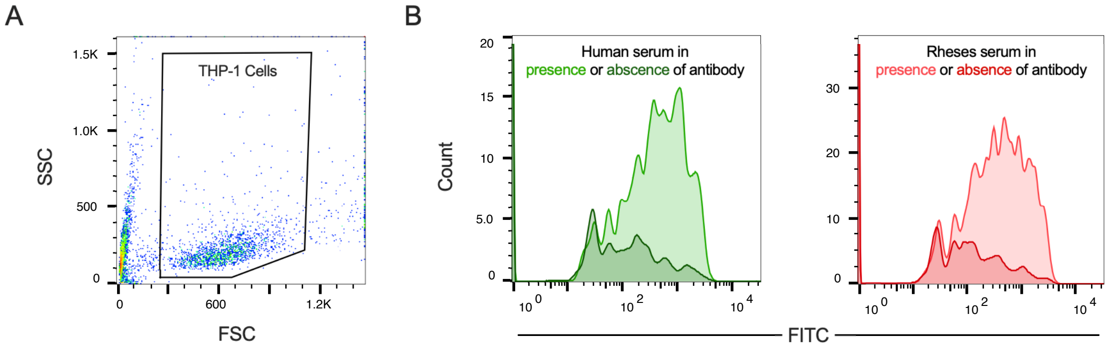


**Supplemental Figure 7: Gating strategy for Complement-aided Antibody-dependent Phagocytosis (C'ADCP). A**. The FSC/SSC gate was drawn to include the THP-1 cells. **B**. Histograms for phagocytosis staining for one of the human serum sample, and one of the rhesus serum sample in presence and absence of bnAb 10-1074.

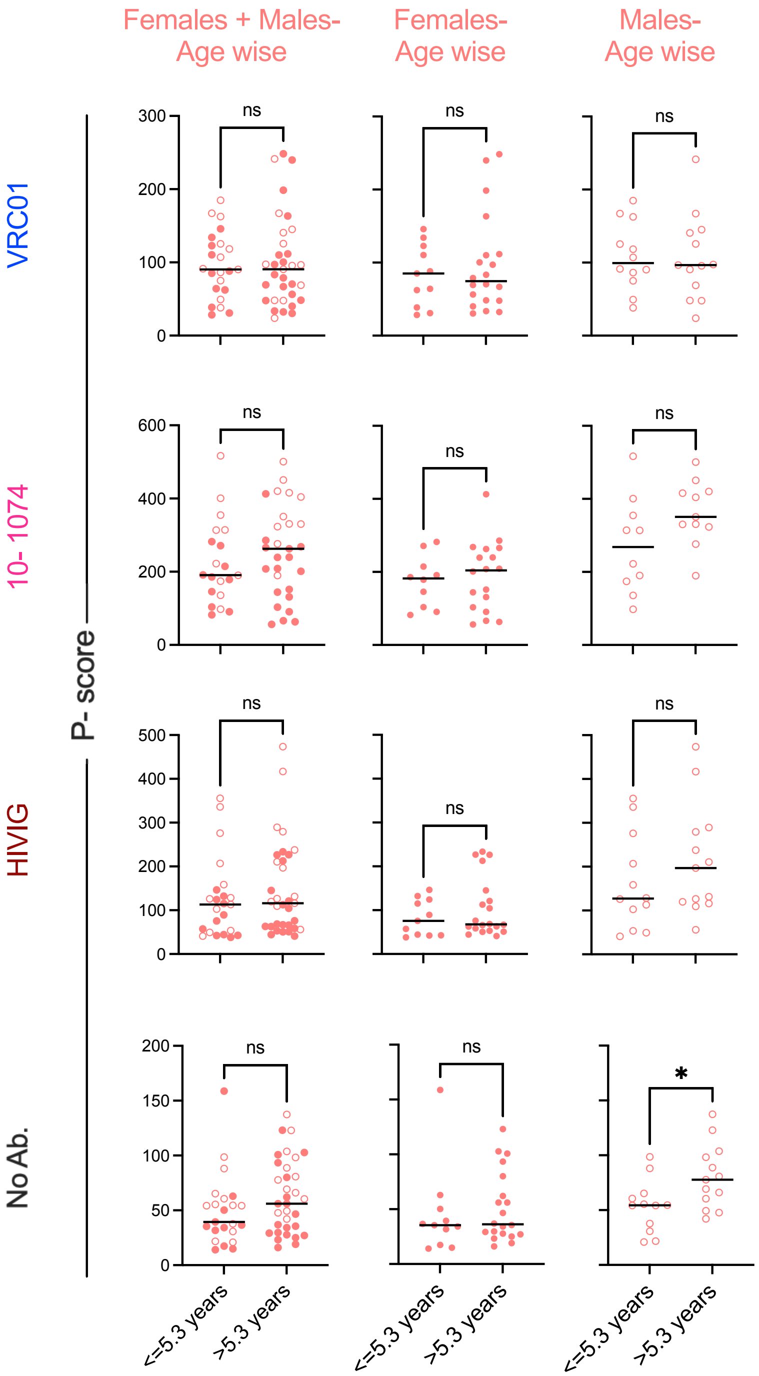


**Supplemental Figure 8: Box plots depicting Complement-aided Antibody-dependent Phagocytosis (C'ADCP) by Rhesus serum.** Box plots of C'ADCP elicited by Rhesus serum based on age of Females+ Males, age of Females and age of Males induced by antibodies VRC01, 10-1074, HIVIG and no antibody control. Statistical significance defined by unpaired t test using Welch's correction (*p≤0.05, ns p>0.05). Bar indicates median.

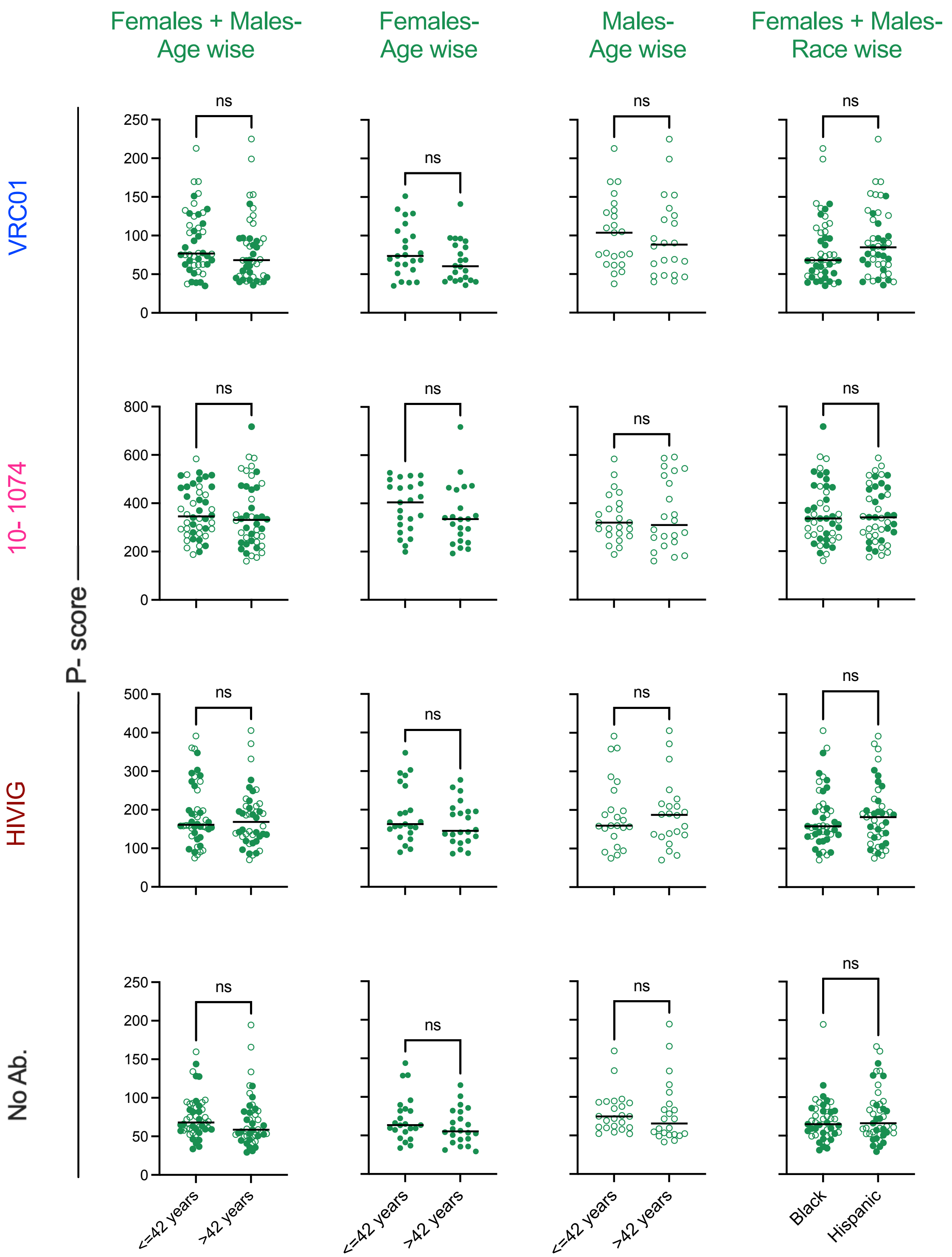


**Supplemental Figure 9: Box plots depicting Complement-aided Antibody-dependent Phagocytosis (C'ADCP) by Human serum.** Box plots of C'ADCP elicited by human serum based on age of Females+ Males, age of Females, age of Males and race induced by antibodies VRC01, 10-1074, HIVIG and no antibody control. Statistical significance defined by unpaired t test using Welch's correction (*p≤0.05, ns p>0.05). Bar indicates median.

**Supplemental Table 1. Study Cohort.** Sample size and donor age in years for human and rhesus samples used in the study.

<table>
<tr><td></td><td colspan="6">Human (n=90)</td><td colspan="6">Rhesus (n=75)</td></tr>
<tr><td>Median age</td><td colspan="6">42</td><td colspan="6">5.3</td></tr>
<tr><td>Average age (s.d.)</td><td colspan="6">43.83 (14.89)</td><td colspan="6">7.15 (4.38)</td></tr>
<tr><td></td><td colspan="3">Females (n=45)</td><td colspan="3">Males (n=45)</td><td colspan="3">Females (n=40)</td><td colspan="3">Males (n=32)</td></tr>
<tr><td>Median age</td><td colspan="3">42</td><td colspan="3">40</td><td colspan="3">6.25</td><td colspan="3">4.95</td></tr>
<tr><td>Average age (s.d.)</td><td colspan="3">44.11 (14.67)</td><td colspan="3">43.56 (15.27)</td><td colspan="3">6.9 (3.46)</td><td colspan="3">7.46 (5.36)</td></tr>
<tr><td></td><td colspan="2">Black (n=45)</td><td colspan="2">Hispanic (n=43)</td><td colspan="2">Caucasian (n=2)</td><td colspan="6"></td></tr>
</table>

**Supplemental Table 2. Antigens used in multiplex assay.**

| Antigen | Catalogue No. | Source |
|---|---|---|
| A.UG37 gp140 | ARP-12063 | NIH HIV Reagent Program, Division of AIDS, NIAID, NIH |
| B.SF162 gp140 | ARP-12026 | |
| B.6240 gp140C | ARP-12572 | |
| B.YU2 gp140 trimer* | ARP-12133 | |
| B.9021 gp140C | ARP-12575 | |
| B.JFRL gp140CF | ARP-12573 | |
| B/C.CN54 gp140 | ARP-12064 | |
| C.1086 gp140C | ARP-12581 | |
| D.UG21 gp140 | ARP-12065 | |
| F.BR29 gp140 | ARP-12066 | |
| M.CON-S gp140 CFI | ARP-12577 | |

*Plasmid was sourced and protein was produced by transient transfection of HEK293 Expi cells (Gibco™, A14635).
